# Ring canals permit extensive cytoplasm sharing among germline cells independent of fusomes in *Drosophila* testes

**DOI:** 10.1101/581702

**Authors:** Ronit S. Kaufman, Kari L. Price, Katelynn M. Mannix, Kathleen Ayers, Andrew M. Hudson, Lynn Cooley

## Abstract

Intercellular bridges, also called ring canals (RCs), connect germline cells during gametogenesis in males and females. They form as a consequence of incomplete cytokinesis during cell division leaving groups of daughter cells connected in syncytia. In *Drosophila* females, RCs are required for oocyte growth but little is known about the role of RCs during spermatogenesis. Using live imaging, we document extensive intercellular movement of GFP and a subset of endogenous proteins through RCs during spermatogenesis from two-cell diploid spermatogonia to clusters of 64 post-meiotic haploid spermatids. Loss of the fusome, a large cytoplasmic structure extending through RCs that is known to be important during oogenesis, has minimal impact on RC development or intercellular protein movement during spermatogenesis. Our results reveal that male germline RCs remain persistently open and mediate extensive sharing of cytoplasmic information, supporting multiple roles for RCs throughout sperm development.

## Introduction

Over millions of years of evolution, gamete development across the animal kingdom has retained the characteristic strategy of maintaining direct cytoplasmic connections between cells during gametogenesis. Intercellular bridges were first discovered by electron microscopy of cat testes in 1955 (Burgos, 1955), and subsequent reports described similar bridges in many other species (Fawcett, 1959; Dym and Fawcett, 1971). The presence of these intercellular bridges (ICBs) is highly conserved from mammals to insects. In mice, ICBs facilitate the intercellular movement of cytoplasm during early oogenesis (Lei and Spradling, 2016). In *Drosophila*, intercellular bridges are called ring canals (RCs) and were first discovered in ovarian cells (Brown and King, 1964; Koch and King, 1966; Koch and King, 1969). In females, RCs are essential for oocyte growth during oogenesis as they allow movement of essential factors required in the transcriptionally silent oocyte (Robinson and Cooley, 1996; Greenbaum et al., 2007). Moreover, previous work from our lab has shown that RCs in *Drosophila* somatic ovarian follicle cells allow movement of cytoplasmic contents between connected cells (Airoldi et al., 2011) that contributes to protein level equilibration (McLean and Cooley, 2013). In contrast, the functional significance of male RCs during spermatogenesis remains less well characterized.

During *Drosophila* spermatogenesis, germline stem cells (GSCs) located at the hub of the testis divide asymmetrically to produce another GSC and a spermatogonial cell that divides mitotically four times to form a cluster of 16 primary spermatocytes connected by RCs (Figure 1A’, B’). Mature spermatocytes enter meiosis synchronously to produce 64 spermatids that remain connected by RCs (Figure 1A”) (Lindsley and Tokuyasu, 1978; Fuller, 1993; Hime et al., 1996). RCs form as the result of incomplete cytokinesis during mitotic and meiotic cell divisions during which cleavage furrows ingress but do not complete the final cytokinetic step of abscission leaving bridges of 1-2 *µ*m in diameter (Figure 1B’). Spermatids remain connected via RCs during the subsequent processes of spermatid tail elongation and individualization that are required for the production of mature, motile spermatozoa (Figure 1B’). Proteins identified at RCs include several that persist from cleavage furrows during cytokinesis as well as proteins recruited after furrow ingression. Unlike the actin-rich intercellular bridges in male mice or *Drosophila* females, *Drosophila* male germline RCs have a septin-rich cytoskeleton, which includes Pnut, Sep1, and Sep2, in addition to Pavarotti (Pav, a kinesin-like protein) and its obligate binding partner Tumbleweed (a RacGAP), the cytoskeletal scaffolding proteins Cindr and Anillin, Nessun dorma, and Orbit/CLASP (for review, see Yamashita, 2018; Haglund et al., 2011).

**Figure 1.**
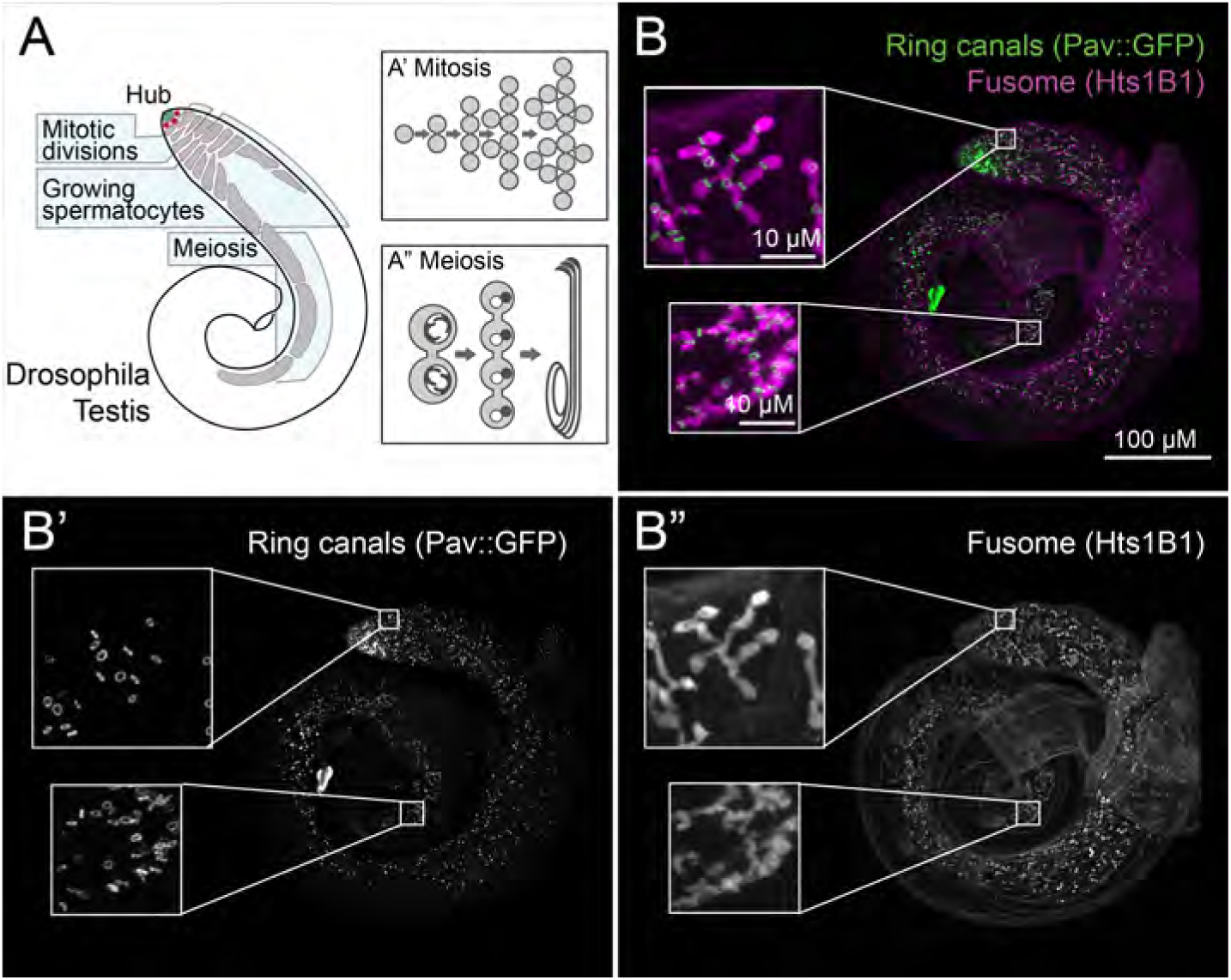
Spermatogenesis overview. (A) Cartoon depicting spermatid development in *Drosophila*. Germline stem cells (red) are located at the hub (green) of the testis. (A’) Spermatogonia divide mitotically four times to form a 16 cell cyst. (A”) These cysts undergo a growth phase of ∼3 days before undergoing two rounds of meiosis to form 64-cell cysts. Following this, each cell elongates a tail to form bundles of mature spermatids. (B-B”) Immunofluorescence shows that ring canals marked by Pav::GFP (B’) and the fusome stained with Hts1B1 (B”) are present throughout spermatogenesis.

In addition to RCs, cells within a cyst are connected by a large, branched organelle that extends through each RC in the cyst (Figure 1B”). Actin, Spectrin, and Adducin (encoded by the *hu-li tai shao (hts)* gene) are all enriched at the fusome (Lin et al., 1994). In females, the fusome is necessary to orient the spindles during cell division and to mediate the transfer of mRNAs and proteins into the pro-oocyte from nurse cells during oocyte specification (Cuevas and Spradling, 1996; Lin et al., 1994). The fusome disappears soon after cells exit the mitotic cell cycle during oogenesis. In males, the fusome persists throughout spermatogenesis until spermatid elongation (Figure 1B”, bottom inset) and has been implicated in the coordinated cell death response of germ cell cysts to DNA damage (Lu and Yamashita, 2017). However, the extent to which the fusome is required for sperm development remains unclear.

Several models have been proposed for the function of germ cell ICBs, all of which invoke the ability of molecules and cytoplasm to move through ICBs between cells; however, evidence for intercellular exchange is limited (Greenbaum et al., 2011). One potential function of ICBs is to allow sharing of cell cycle signals required for synchronous cell divisions like those seen in both *Drosophila* and mammalian testes (Fuller, 1993; Huckins, 1978; Ren and Russell, 1991; Gärtner et al., 2014). A second hypothesis for ICB function is in quality control as elimination of a single defective cell might initiate propagation of cell death signals through ICBs to the rest of the cells to efficiently eliminate the compromised cyst (Greenbaum et al., 2011; LeGrand, 2001). A recent study in the *Drosophila* testis revealed that loss of connectivity between the cells mediated by RCs or the fusome may reduce movement of a cell death marker between the cells (Lu and Yamashita, 2017). A third possible function of ICBs is to allow movement of X-linked gene products into Y-bearing spermatids and vice versa. As a result, haploid cells could remain phenotypically diploid by inheriting products from both X- and Y-chromosomes (Fawcett, 1959; Erickson, 1973; Braun et al., 1989). This idea is supported by several key studies. In *Drosophila*, Lindsley and Grell (1969) showed that functional sperm are produced despite lacking either one or both major autosomes. A study in mouse demonstrated that mRNA and protein from a single copy transgene were distributed evenly in all haploid spermatids (Braun et al., 1989). Additional evidence from mouse has shown that organelles and protein complexes also pass through post-meiotic ICBs (Morales et al., 2002; Ventelä et al., 2003). While evidence supports intercellular cytoplasm mobility during spermatogenesis in both mice and flies, systematic examination of intercellular movement through ICBs or RCs has not been reported.

Here, using a combination of extensive live cell imaging, genetics, and electron microscopy, we explored the function of *Drosophila* male germline RCs during spermatogenesis. We directly observed movement of GFP and endogenous proteins through RCs. Movement between neighboring cells within a germline cyst occurred throughout all stages of spermatogenesis, including post-meiotically, and independently of protein size. Further, we found that the fusome did not have a major role in RC formation or intercellular protein movement.

## Results

### RCs allow intercellular movement of proteins in mitotic spermatogonial cells

To investigate intercellular protein movement in the *Drosophila* testis, we expressed photoactivatable GFP (PA-GFP) (Pfeiffer et al., 2012) using *nos-* or *bam-Gal4* drivers in germline cells also expressing GFP-tagged Pav (Pav::GFP) to mark RCs. Following activation in a single cell within a spermatogonial cyst, we captured time-lapse movies of PA-GFP localization (Figure 2, Figure S7-S10). Within 30 seconds following photoactivation, we observed movement of PA-GFP from the activated cell through RCs to other cells within mitotically active cysts (Figure 2A-L). Within 10 minutes following photoactivation, we observed GFP signal throughout most, if not all, cells in all spermatogonial cysts from 2-cell through 16-cell cysts (Figure 2M-P, n=94). These data demonstrate that RCs are persistent open channels that allow diffusion of GFP between spermatogonial cells.

**Figure 2.**
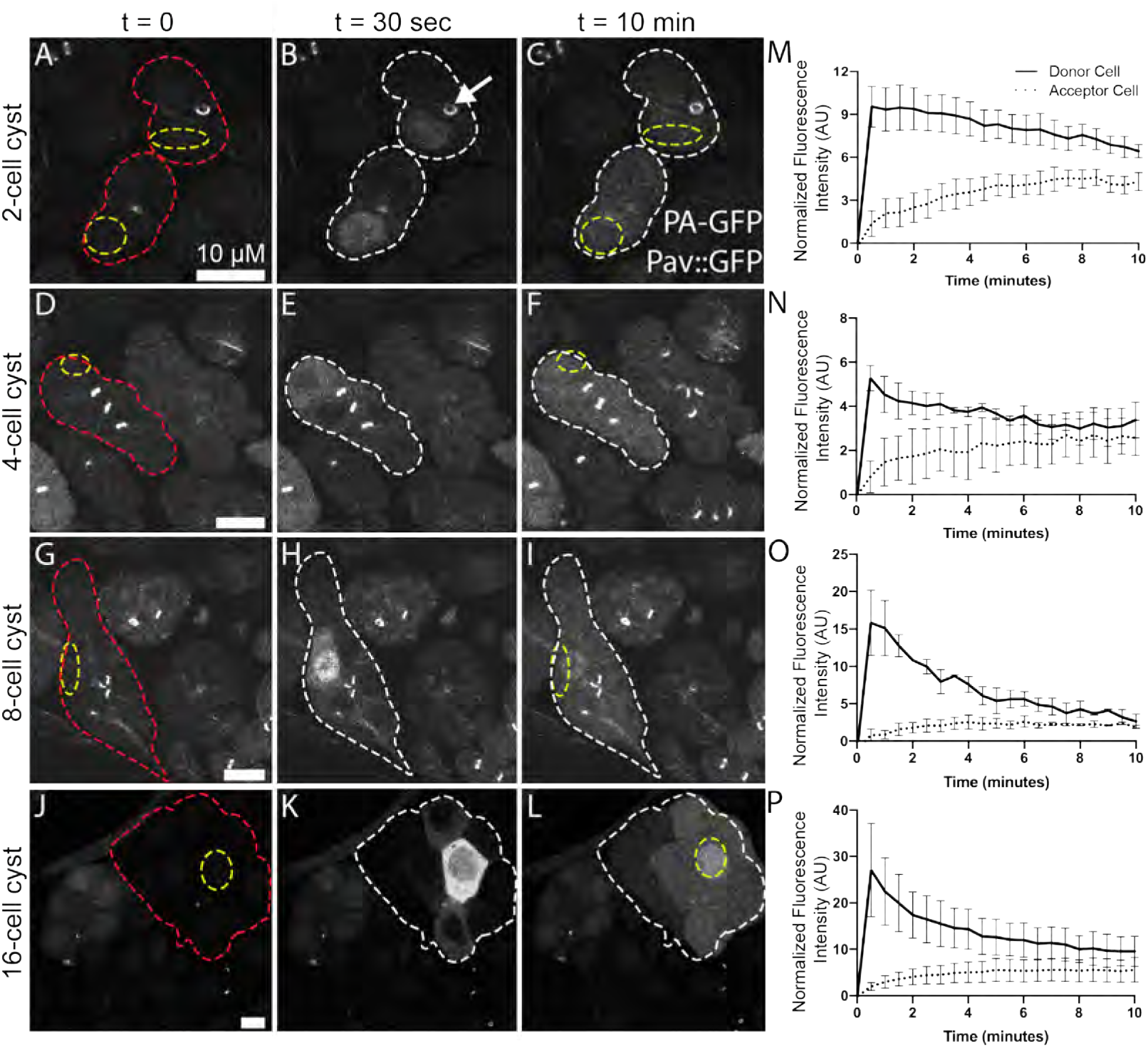
RCs allow movement of GFP between cells in a cyst. (A-L) Live imaging of activated PA-GFP at various stages of spermatogenesis reveals extensive sharing of GFP between cells in a cyst (red outline) through the ring canals (marked with Pav::GFP, white arrow). After activation of PA-GFP in a single cell or small region of cells (yellow outline), GFP was found in most of the cells in that cyst after 10 minutes (white outline). (M-P) Quantification of PA-GFP movement following photoactivation from a single donor cell (solid line) to other cells within the cyst (dashed line). Relative fluorescence intensity (AU) was plotted with respect to time. Error bars represent SEM.

To examine the movement of endogenous *Drosophila* proteins through RCs, we performed Fluorescence Loss in Photobleaching (FLIP) experiments in 16-cell post-mitotic spermatocyte cysts expressing either GFP or GFP-fusion proteins. We selected FlyTrap lines with GFP-tagged proteins (Buszczak et al., 2007; Lowe et al., 2014; Nagarkar-Jaiswal et al., 2015; Quiñones-Coello et al., 2007) that had high GFP-expression in the male germline (Figure S1) or had been previously assayed for movement in female somatic follicle cells (Airoldi et al., 2011). A region of interest was repeatedly photobleached while we simultaneously captured images of the GFP fluorescence in the entire cyst. Movement of proteins into the bleaching zone was determined by a loss of GFP fluorescence in neighboring cells. We first demonstrated by FLIP that, consistent with PA-GFP results, cytoplasmically expressed GFP moved through the RCs (Figure 3A-C, M). Similarly, we observed movement of several GFP-fusion proteins including Kra, Oda, Men-B, and *β*Tub56D (Figure 3D-I, Table 1). However, we did not detect movement of most of the selected GFP-fusion proteins within the 60 minute time-frame (Table 1). The size of the different proteins did not appear to correlate with the ability to move through the RCs between the cells. Remarkably, GFP::CaM, which we previously showed moves through RCs in female follicle cells (Airoldi et al., 2011), did not move between male germline cells (Figure 3J-L, P). These data indicate that only a subset of proteins freely diffuses between cells despite the open intercellular bridges.

**Table 1.** RCs allow sharing of some, but not all, proteins. Fluorescence Loss in Photobleaching (FLIP) demonstrates that some, but not all, GFP-tagged proteins move between the cells in a 16-cell cyst.

| Protein | Size (kDa) | Function | FLIP? |
| --- | --- | --- | --- |
| GFP | 27 | Fluorescent protein | Yes |
| Kra | 49 | Translational regulator | Yes |
| Men-B | 69 | Metabolism enzyme | Yes |
| Oda | 28 | Antizyme | Yes |
| βTub56D | 51 | Cytoskeletal constituent | Yes |
| CaM | 17 | Calcium-binding messenger protein | No |
| Clu | 161 | Cytoplasmic ribonucleoprotein | No |
| CG32701 | 33 | Unknown | No |
| eIF4A | 46 | Eukaryotic translation initiation factor | No |
| eIF4E1 | 46 | Eukaryotic translation initiation factor | No |
| Lost | 60 | RNA metabolism protein | No |
| Mito-GFP | - | Mitochondrial targeting sequence | No |
| Pdcd4 | 56 | Stem cell differentiation | No |
| Sgg | 59 | Meiotic kinase | No |

**Figure 3.**
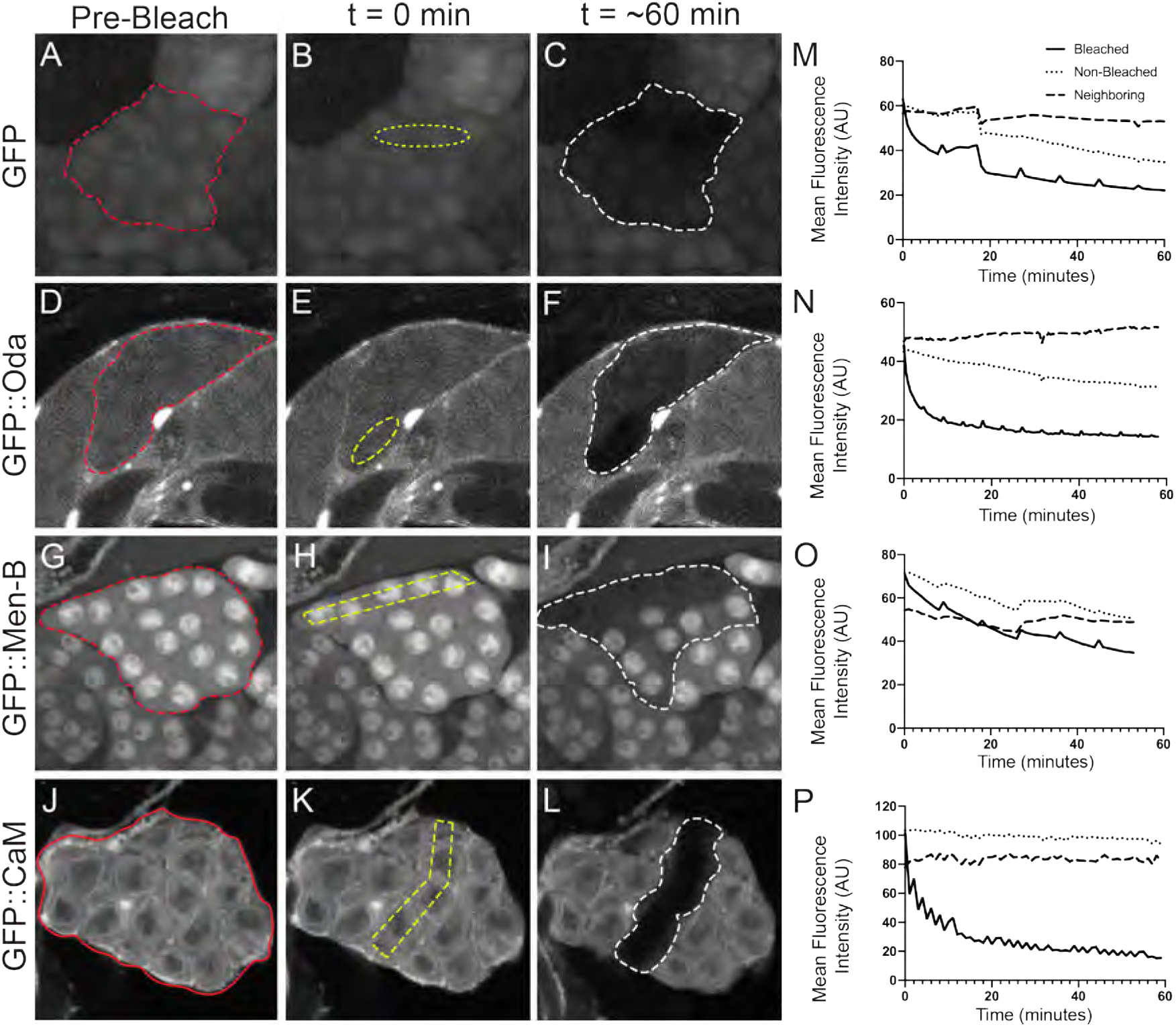
RCs allow for sharing of some, but not all, proteins. (A-L) Fluorescence Loss in Photobleaching (FLIP) demonstrated that some, but not all, GFP-tagged proteins move between the cells in a 16-cell cyst. Several cells within a 16-cell cyst (red outline) expressing GFP or a GFP-tagged protein were continuously bleached (yellow outline) over the course of ∼1 hour. Movement of a protein was determined by a loss in GFP fluorescence from neighboring cells within that cyst (white outline) indicating that GFP from non-bleached cells moved into the bleach region. Movement of cytoplasmic GFP, GFP::Oda and GFP::Men-B was detected, while GFP::CaM did not move between the cells. (M-P) Quantification of GFP in the bleached (solid line), non-bleached (dotted line), and neighboring (dashed line) regions in a spermatocyte cyst. FLIP was detected for GFP, GFP::Oda, GFP::Men-B but not GFP::CaM. Mean fluorescence intensity (AU) is plotted with respect to time.

### Fusome disruption has minimal impact on male fertility

During *Drosophila* oogenesis, the fusome is necessary for the production of viable gametes (Cuevas and Spradling, 1996; Lin et al., 1994). To investigate the role of fusomes during spermatogenesis, we carried out germline-specific RNAi of *α-Spectrin* by driving *α-Spectrin* shRNA with *nos-* or *bam-Gal4*. In wild-type testes, the fusome extended through the RCs connecting all the cells within a cyst (Figure 4A, B). In *α-Spectrin* RNAi testes, we detected no Spectrin (Figure S2C, D) or Hts (Figure 4C, D) fusome labeling, indicating the fusome was either compromised or eliminated. Surprisingly, overall testis morphology appeared normal with all stages of spermatogenesis present (Figure S3). Furthermore, fusome disruption had negligible effect on male fertility. *α-Spectrin* RNAi males displayed a slight increase in fertility and knockdown of *hts* by RNAi caused no observable change in fertility (Figure S4A). In contrast, several loss-of-function alleles of *hts* displayed marked reductions in male fertility (Figure S4B, C), as previously shown by Wilson (2005), suggesting a requirement for *hts* in non-germline cells for fertility.

**Figure 4.**
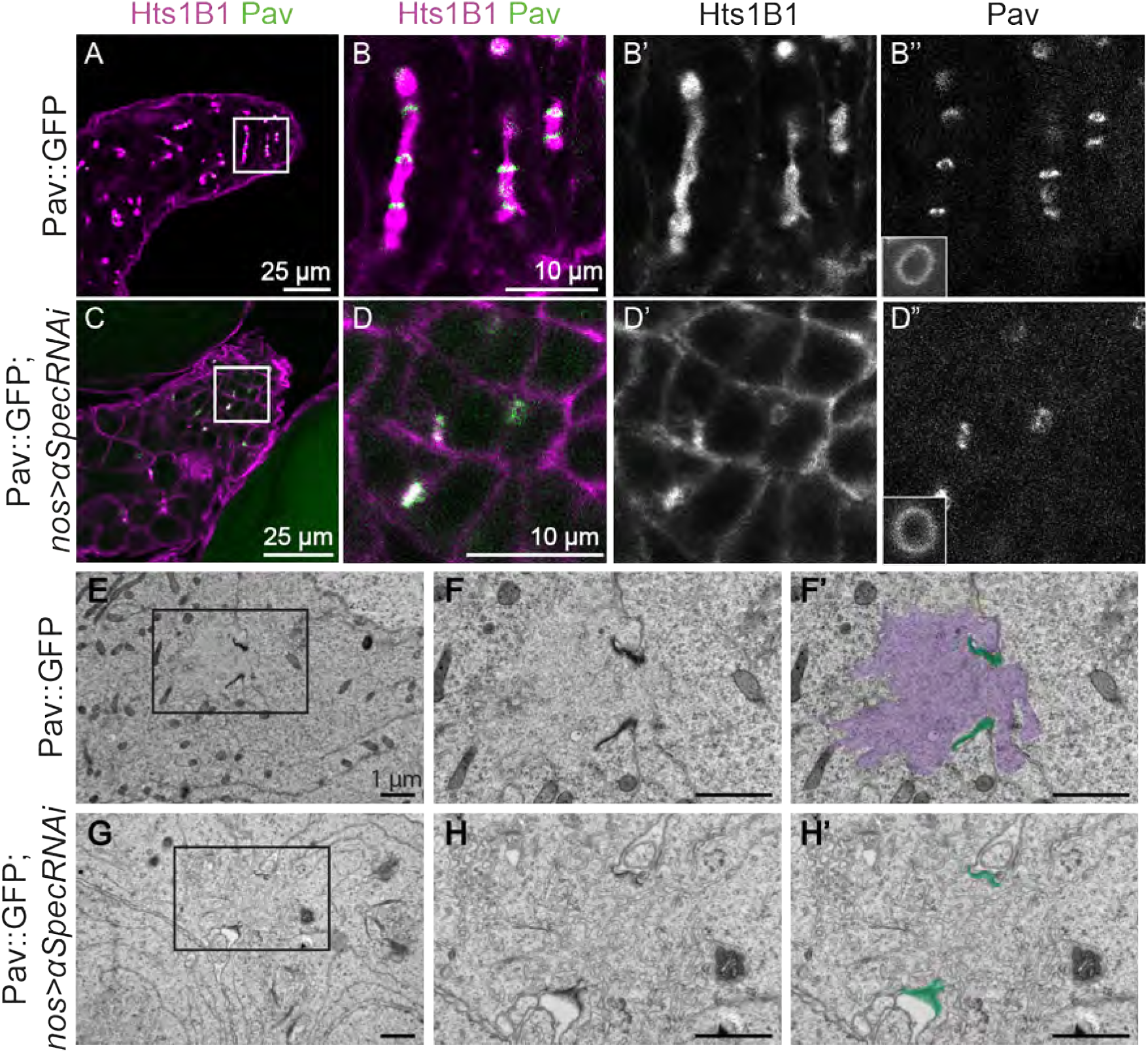
Fusome loss does not have effect on cyst development. (A) Wild-type testis with RCs marked by Pav::GFP and fusomes labeled with Hts1B1 antibody. (B-B”) The region marked by box in (A), highlighting the RCs (Pav, green) and fusome (Hts1B1, purple) in wild-type testis. Inset in B” highlights one RC. (C) Testes with *α-Spectrin* RNAi lack Hts1B1 staining at the fusome, but testis morphology is unaffected. (D-D”) The region marked in (C) showing Hts1B1 staining at the membrane rather than in a fusome pattern while Pav::GFP remained localized to the RCs. (E) EM of Pav::GFP testis revealed electron dense RCs surrounding a fusome. (F) Inset from (E) shows electron dense RCs (green in F’) and a ribosome-free fusome area (purple in F’). (G) EM of two cells connected by a RC in a *α-Spectrin* RNAi testis. (H) The region marked by a box in G. RCs were still present, though they appeared less electron dense. Importantly, no fusome was distinguishable from the rest of the cytoplasm. (H’) False coloring of (H) highlighting RC presence (green) but lack of clearly marked fusome. NOTE: A PDF FILE WAS SUBMITTED FOR VIEWING HIGH RESOLUTION OF EM IMAGES.

Our data suggested that fusomes are not required for male fertility. However, it was possible that removing the Adducin (encoded by *hts*) and *α*-Spectrin proteins only affected the fusome cytoskeleton but did not completely disrupt the fusome. To investigate fusomes in *nos-Gal4* > *α-Spectrin*^*RNAi*^ testes at higher resolution, we used electron microscopy (EM). In wild-type testes, cross-sections of RCs were easily identified by their electron-dense membranes (n=21, Figure 4E-F’, Figure S5A, B). The fusome (Figure 4F’, purple shading) appeared as a cloud of ribosome-free cytoplasm extending through the RCs (Figure 4F’, green shading) and between connecting cells. In both *nos-Gal4*> *α-Spectrin*^*RNAi*^ (n=7) (Figure 4G) and *hts* loss-of-function mutant (n=5) (Figure S5 C-F) samples, the RC backbones appeared somewhat less dense. More importantly, there were no obvious ribosome-exclusion zones indicating that the fusomes were completely missing (Figure 4H-H’, Figure S5 C-F). Thus, fusomes are dispensible for male fertility.

Although RCs appeared somewhat less dense in EMs, we found that fusome loss did not affect accumulation of Pav::GFP to RCs (compare Figures 4B” and 4D”). Furthermore, the diameters of the RCs in wild-type and fusome-less testes were the same at 1.6 *µ*m (n=300 wild-type and 201 *nos* > *α-Spectrin*^*RNAi*^ RCs, p=0.98). However, 8.3% of RCs in the fusome-less testes appeared morphologically abnormal or collapsed, suggesting that the fusome may play a role in RC stability. Given that the germline-specific fusome knockdown testes appeared morphologically normal overall, RCs were present as evidenced by Pav::GFP localization, and males were fertile, we conclude that fusomes are not necessary for male fertility under normal conditions. This is in stark contrast to Drosophila females where loss of fusomes causes oogenesis arrest and sterility (for review see Huynh (2013)).

### The fusome is not necessary for movement of GFP between cells

We explored whether disruption of the fusome had an impact on protein movement through the RCs during spermatogenesis by repeating our PA-GFP analysis in *α-Spectrin* RNAi testes (Figure 5, Figure S14-S15). After photoactivation in a single cell, PA-GFP moved to neighboring cells in 2-, 4-, and 16-cell cysts, just as in wild-type testes (n=93). Moreover, FLIP analysis in a fusome knockdown background showed that, as shown in wild-type testes, GFP::Oda did move between cells whereas GFP::CaM did not (data not shown). Therefore, the fusome is not necessary to mediate transport between the cells in a cyst.

**Figure 5.**
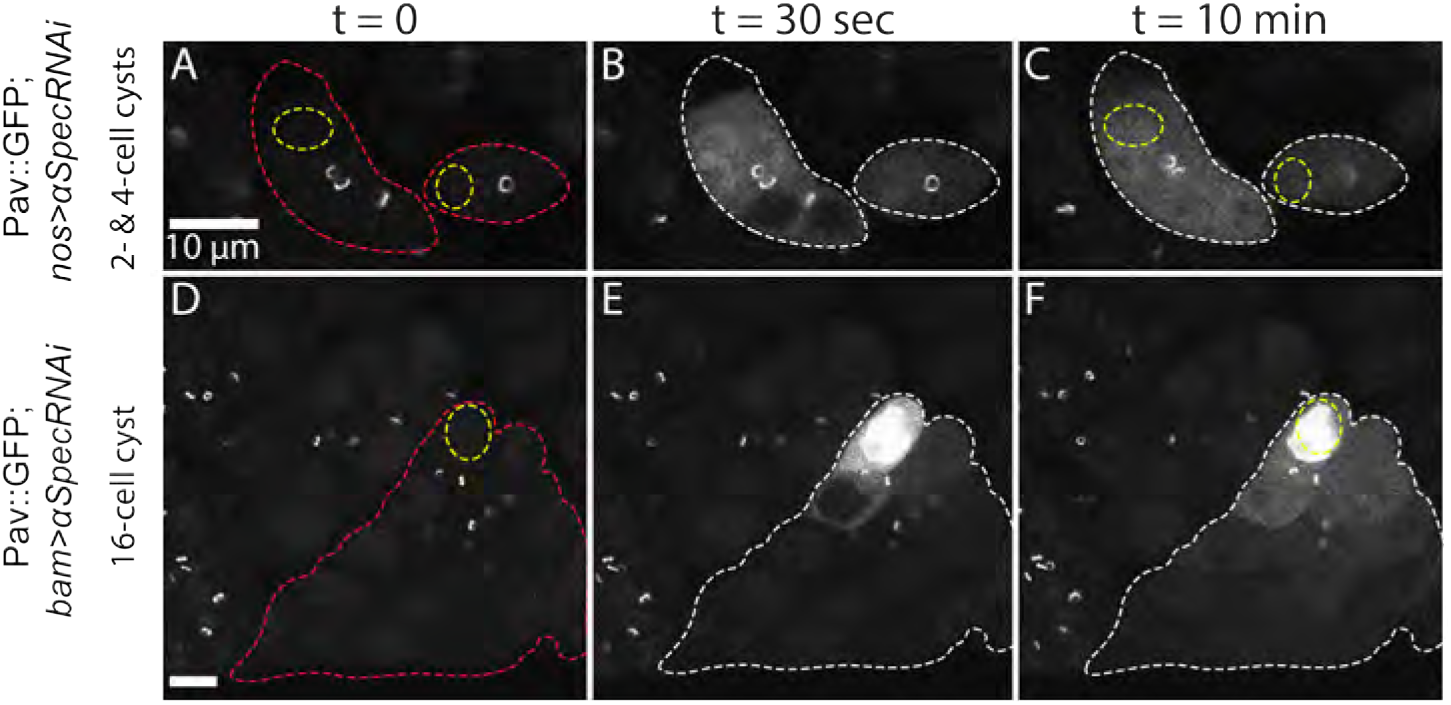
PA-GFP moves through RCs in the absence of fusome components. *α-Spectrin* RNAi driven by *nos-Gal4* in 2- and 4-cell (A-C) and 16-cell (D-F) spermatocytes cysts (red outlines) did not affect movement of GFP through RCs (marked with Pav::GFP). PA-GFP was activated in one cell (yellow outline) and moved through the RCs to other cells within that cyst (white outline).

To determine if the fusome regulates the rate of exchange between cells, we needed to capture both higher time resolution of movement between cells and the total fluorescence in the entirety of the cells. To accomplish these goals, we used swept field microscopy to capture z-stacks of spermatocyte cysts. We activated PA-GFP in a single spermatocyte cell and tracked GFP movement in a z-projection encompassing the volume of the cyst. We focused on cysts where PA-GFP diffused from the activated cell (Figure 6A, yellow outline) into a single adjacent recipient cell (Figure 6A, white outline), connected by a RC (Figure 6A, arrow). We measured the total fluorescence of PA-GFP in the activated and recipient cells and plotted the mean relative fluorescence units (RFU) of the recipient cell over time (Figure 6B; wild-type n=9, *bam* > *α-Spectrin*^*RNAi*^ n=11). We plotted a non-linear regression of PA-GFP RFU mean values (Figure 6B, dashed lines) and fitted curves (Figure 6B, solid lines) for both wild-type (blue lines) and fusome RNAi (red lines) testes. The fitted curves closely approximate the mean RFU values (wild-type R^2^ = 0.95, *bam* > *α-Spectrin*^*RNAi*^ R^2^ = 0.83). Although the RFU plateaus of the curves were not significantly different, the diffusion rate (K) was significantly higher when the fusome was absent (p<0.0001) (Figure 6C). Also, there was more variability in the rate of movement upon fusome knockdown (Figure 6B). These data suggest that while both cells ultimately accumulate comparable levels of GFP, the rate of accumulation is different between the genotypes; the fusome-less cells have a faster rate of diffusion (.008947 RFU/s) compared to the wild-type controls (0.007343 RFU/s). Taken together, these results demonstrate that the fusome is required for proper protein movement dynamics and suggest that the fusome may function as a diffusion barrier between wild-type cells.

**Figure 6.**
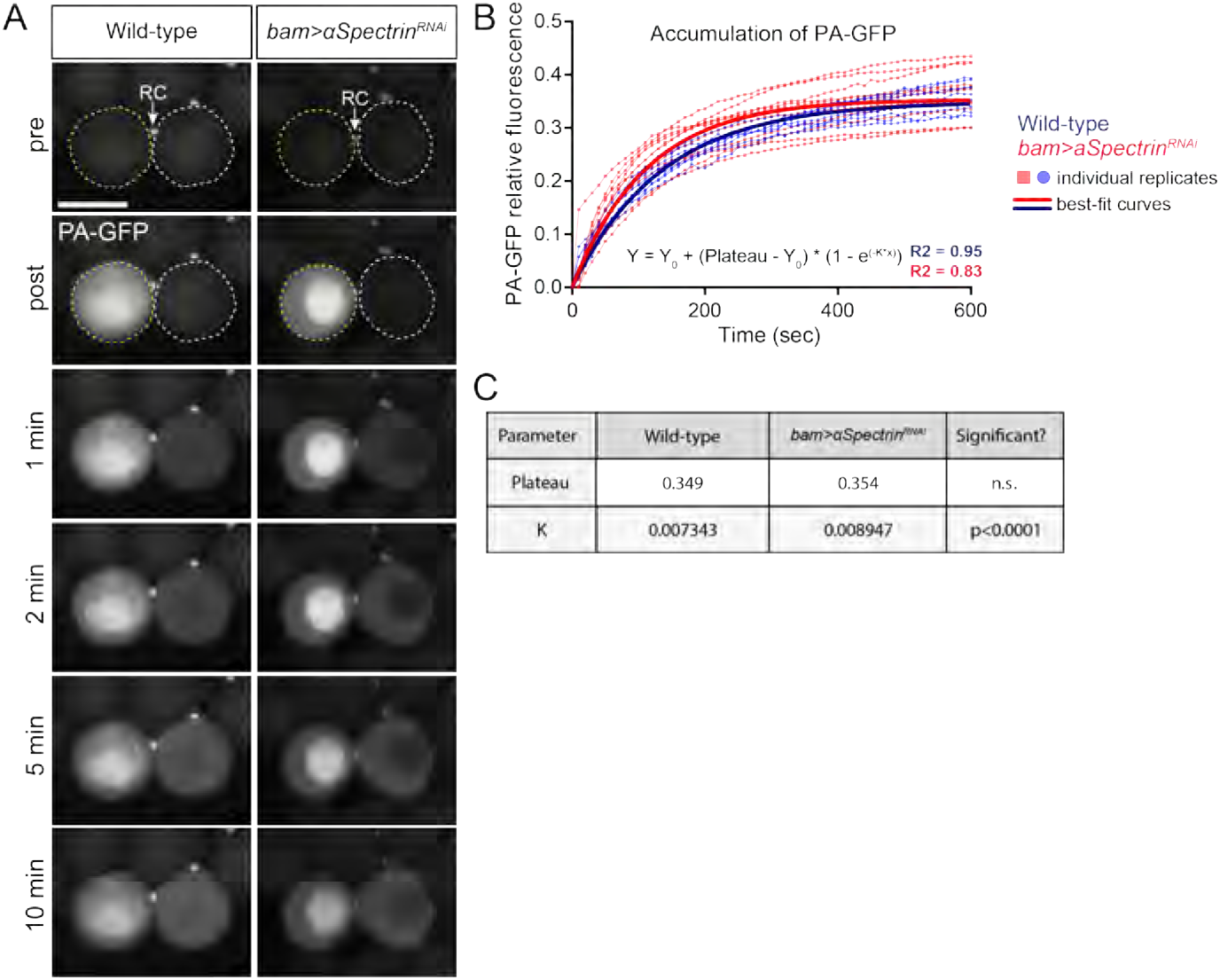
Movement of PA-GFP through RCs in fusome knockdown cells is faster than in wild-type cells. (A) Single cells expressing PA-GFP within a spermatocyte cyst were activated (yellow dashed outline) and PA-GFP fluorescence between the activated and recipient cell (white dashed outline) was imaged throughout a 10 minute time course in wild-type and fusome knockdown testes. Arrows indicate the RC, marked by Pav::GFP. Two-cell systems were selected for quantification, meaning that only movies in which PA-GFP diffused into a single adjacent cell were used for analysis. (B-C) Non-linear regression of PA-GFP RFU mean values (dashed lines) and fitted curves (solid lines) for both wild-type (blue lines) and fusome RNAi (red lines) testes. Both curves plateaued at the same RFU, but the rate of movement (as measured by the K rate constant parameter) in fusome RNAi was significantly faster than in wild-type (p<0.0001). wild-type: n=9; fusome RNAi: n=11. Scale bar = 20 *µ*m.

### Proteins can move between cells in meiotic and post-meiotic cysts

One hypothesis for the function of RCs during spermatogenesis is to allow sharing of X-linked gene products to Y-bearing cells after meiosis (Braun et al., 1989; Morales et al., 2002). To test this idea, we performed photoactivation studies in meiotic and post-meiotic cysts as well as haploid spermatids. In 32- and 64-cell cysts, we could easily detect the spread of photo-activated PA-GFP between cells (Figure 7A-C,D-F, Figure S11-S12). We observed GFP fluorescence in all cells of a 32-cell cyst within 10 minutes following photoactivation (Figure 7A-C, Figure S11). It took more time for all cells within a 64-cell cyst to accumulate enough GFP fluorescence to visualize (Figure 7D-F, Figure S12). Movement of PA-GFP through the RCs in post-meiotic cells was not impacted by the loss of fusome, as PA-GFP still moved across RCs in the spermatid stage in the *α-Spectrin* RNAi testis (Figure 7G-I, Figure S16). Additionally, we performed FLIP on endogenous GFP-tagged proteins in 64-cell cysts, and found their ability to move between cells the same as we observed in primary spermatocyes (GFP::Men-B shown in Figure 7M-O).

**Figure 7.**
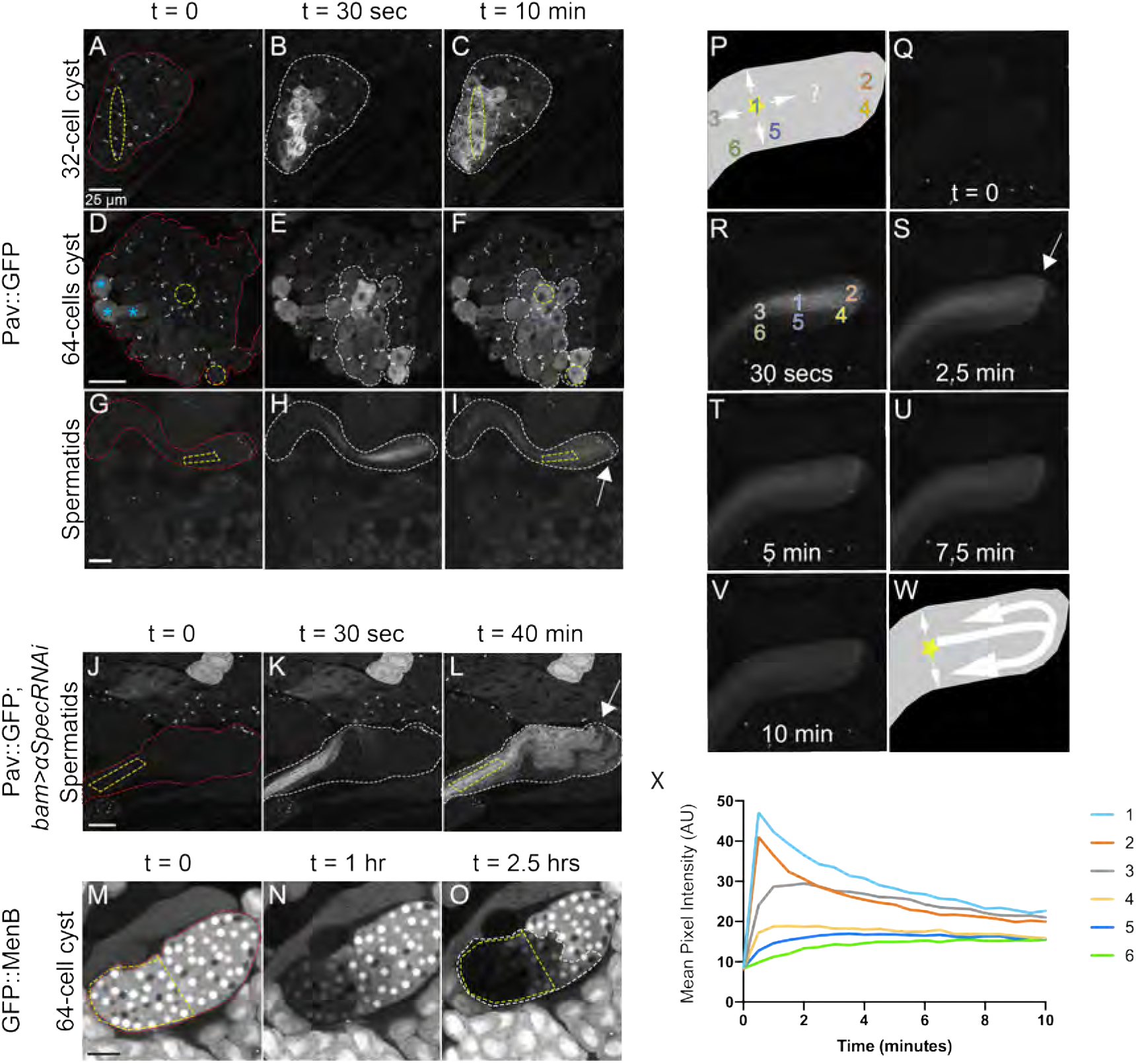
Movement of proteins in meiotic cysts and haploid spermatids. (A-I) After meiosis I and II, and during elongation of spermatid tails, PA-GFP moved between cells of a cyst (red outline). PA-GFP activated within a small region of the cyst (yellow outline) appeared in neighboring cells within that cyst (white outline). Cells activated previously in (D) are marked with blue asterisks. (J-L) PA-GFP movement occurred through RCs in elongated spermatids even after loss of the fusome with *α-Spectrin* RNAi. (M-O) Movement of endogenous GFP::Men-B occurred in post-meiotic 64-cell cysts. Bleaching zone in this FLIP experiment is outlined in yellow. Cells with loss of fluorescence are outlined in white. (P) Cartoon of spermatid bundle depicting possible pathways of PA-GFP spread after activation (marked by star at position 1). (Q-V) PA-GFP activation in a spermatid bundle over the course of 10 minutes with activation occurring in region 1 (shown in panel R). (W) Cartoon of actual PA-GFP spread showing that movement is predominantly through RCs rather than through lateral perforations. (X) Normalized fluorescence intensity of PA-GFP over time measured at the regions indicated in (R). GFP fluorescence increased in region 4 (yellow line) before region 5 (dark blue line). RC end of spermatid bundles marked by arrow.

Finally, we tracked movement of PA-GFP in elongated spermatids after activation in a central zone of a spermatid bundle. We observed the spreading of GFP along the length of the bundle and to outer spermatids (Figure 7G-I, Figure S13). Since RCs were located at one end of the spermatid bundle (Figure 7I, arrow), GFP would need to reach RCs before spreading to lateral cells. However, in addition to the RCs, elongated spermatids contain lateral membrane perforations throughout the length of the tails (Tokuyasu et al., 1972; Fabrizio et al., 1998), making it possible that GFP moved laterally through the perforations independent of RCs (Figure 7P). To investigate protein movement further, we carefully tracked the spread of PA-GFP over a period of 10 minutes following activation (Figure 7Q-V). We reasoned that if movement to lateral cells depends on RCs, GFP fluorescence would increase near RCs (Figure 7P, region 4) before it accumulated at a location directly lateral to the activation zone (Figure 7P, region 5). In all samples examined (n=3), PA-GFP was first observed in region 4 (Figure 7X, Figure S6). Although we cannot rule out some lateral movement through perforations, these data suggest that intercellular movement is predominantly through RCs (Figure 7W). Our observation of intercellular protein movement during and after meiosis and during spermatid tail elongation supports the hypothesis that X-linked proteins can be shared between haploid cells.

## Discussion

Although intercellular bridges connect germline cells in testes throughout the animal kingdom, their role during spermatogenesis has remained unclear. Here we establish for the first time that RCs in *Drosophila* testes remain open throughout spermatogenesis from spermatogonial through elongated spermatid stages. Using live imaging approaches, we documented diffusion of GFP and endogenous proteins through RCs in all stages, including post-meiotically. Ring canals are located at the opposite end of the nuclei in haploid spermatid bundles; we observed rapid GFP diffusion along the length of bundles, and spreading to neighboring cells by way of RCs. In addition to RCs, *Drosophila* germline cells contain fusomes, which provide another type of connectivity between cells. We disrupted fusomes by RNAi inhibition of structural components and found negligible effects on fertility. Furthermore, movement of GFP between cells lacking fusomes was changed only in the rate of exchange, suggesting that fusomes are not needed to actively promote or prevent protein movement through RCs. Our work represents the first extensive evidence that RCs mediate the sharing of cytoplasmic components throughout the entire process of *Drosophila* spermatogenesis. These data support several possible roles for RCs during spermatogenesis including sharing X-linked gene products among haploid cells, involvement in cell cycle synchronization, and promotion of overall cyst health.

We detected movement of some, but not all, proteins through RCs during spermatogenesis within the time frame of our live imaging experiments. The size of a protein does not appear to correlate with its ability to move into neighboring cells. Proteins ranging from 28 to 69 kDa moved through RCs, while many proteins in the same range of molecular mass did not (Table 1). These results are similar to our assessment of movement across the smaller RCs connecting somatic follicle cells in *Drosophila* ovaries (Airoldi et al., 2011). An unexpected difference is that GFP-tagged Calmodulin (CaM), a 17 kDa calcium binding messenger protein, moved freely through follicle cell RCs, but not through RCs in the testis. Previous work has shown differences in CaM diffusion rates depending on whether CaM is bound to other complexes or immobile structures (Sanabria et al., 2008). The ability of proteins to diffuse between cells may be dependent on whether they are associated with a larger complex, an organelle, or the cytoskeleton.

Our work supports the long-held idea that male RCs are required to allow movement of gene products between post-meiotic cells so that haploid cells remain phenotypically diploid. While active biosynthesis of crucial mRNAs and proteins occurs during the 3-day growth phase of the primary spermatocytes prior to meiosis (Fuller, 1993), a subset of genes is transcribed only after meiosis during spermatid tail elongation, called “cup” and “comet” genes based on the localization of mRNAs clustered at one end of spermatid bundles (Barreau et al., 2008; Wilk et al., 2017). The timing of cup and comet gene expression suggests that they function during spermatid development. Mutants of one comet gene, *scotti*, which is involved in spermatid individualization, are male sterile (Barreau et al., 2008). Since *scotti* heterozygotes are fertile, products made in haploid spermatids with the wild-type allele likely spread to spermatids with the mutant allele (White-Cooper, 2010). Although this equilibration could be mediated by lateral perforations between spermatid tails (Tokuyasu et al., 1972; Fabrizio et al., 1998), our work suggests the most efficient path between cells is indeed through RCs. This would be especially critical for products of post-meiotically expressed X-linked genes such as *r-cup* and *p-cup*.

Discovering that fusomes are virtually dispensable in *Drosophila* testes was unexpected. Disruption of fusomes in ovaries causes egg chamber arrest and female sterility (Lin et al., 1994; Yan et al., 2014), while we found that germline RNAi of fusomes in males has no measurable effect on male fertility. The composition of fusomes is also different in males and females; while they both have the same cytoskeletal proteins (*α*-Spectrin, Adducin, Ankyrin), only female fusomes are rich in endoplasmic reticulum (ER) membranes and ER proteins (Lighthouse et al., 2008; Yamashita, 2018; Snapp et al., 2004; Cuevas et al., 1997; Hime et al., 1996). Interestingly, while male fusomes persist (Hime et al., 1996), female fusomes are present only in the oogonial stage of development where they have been implicated in mitotic spindle orientation, cell cycle control and oocyte specification (Yamashita, 2018; Lin et al., 1994; Lilly et al., 2000; McGrail and Hays, 1997; Cuevas and Spradling, 1996; Deng and Lin, 1997; Lin and Spradling, 1995). Fusome disassembly then happens as a polarized microtubule cytoskeleton forms that allows the single oocyte to be nourished by its 15 sibling nurse cells (Grieder et al., 2000). In contrast, all 16 cells of male cysts produce sperm so a structural mechanism is not needed for an asymmetric cell fate specification. While there is evidence that male fusomes participate in mitotic spindle alignment (Miyauchi et al., 2013), we can now conclude that fusomes are not essential during either mitosis nor meiosis during spermatogenesis under normal conditions. It was recently reported that, as a response to DNA damage, the male fusome facilitates the movement of a cell death marker through RCs between connected cells during spermatogenesis and that this is necessary for proper sperm production (Lu and Yamashita, 2017). Perhaps RCs, and the persistence of the fusome extending through them, are retained in males to ensure proper quality control during spermatogenesis during times of stress.

Our live imaging data support a function for RCs during spermatogenesis in mediating extensive cytoplasmic sharing between the cells in a cyst; however, more could be learned about RC function through their targeted disruption or occlusion. Attempts at RC disruption have focused on RNAi of RC proteins but these efforts led to cytokinesis defects rather than a RC-specific phenotype. To progress, new tools must be developed using innovative strategies to disrupt RC function *in vivo*, perhaps by occlusion or targeted disruption, to study the functional consequences of RC loss during spermatogenesis. Disruption of the RCs in this manner could provide additional evidence for RC involvement in cell-cycle synchronization, maintenance of overall cyst health, and sharing of post-meiotic gene products.

## Materials and Methods

### *Drosophila* strains

The following *Drosophila* lines were generously provided by the referenced authors: 20XUAS-mC3PA-GFP and 10XUAS-GFP (Pfeiffer et al., 2012), *bam-Gal4* (Chen and McKearin, 2003), *hts*^*1*^ (Yue and Spradling, 1992), *hts*^*W532X*^ (gift from Trudy Schüpbach), and *hts*^Δ*G*^ (Koundakjian et al., 2004). The following FlyTrap lines were used: GFP::CaM (YC0069LE), eIF*α*::GFP (YC0001), and GFP::Oda (YD0523) (Buszczak et al., 2007; Lowe et al., 2014; Nagarkar-Jaiswal et al., 2015; Quiñones-Coello et al., 2007). *nos-Gal4* (stock #7303), *αSpectrin* shRNA (stock #56932), *hts*^*1103*^ (stock#10989), *hts* shRNA (stock #35421), Df(2R)BSC135/CyO (stock #9423), GFP::Clu (stock #6842), GFP::eIF4E1 (stock #50858), GFP::CG32701 (stock #50839), GFP::Lost (stock #6832), Mito::GFP (stock #8442), Pdcd4::GFP (stock #38446), GFP::Sgg (stock #50887), GFP::Kra (stock #50873), GFP::Men-B (stock #50854), and GFP::*β*Tub56D (stock #50867) were obtained from the Bloomington Drosophila Stock Center.

### Construction of transgenes and generation of transgenic lines

To visualize RC components at endogenous levels, we recombineered GFP into a BAC containing the Pavarotti (Pav) gene at the C-terminus (BAC ID 322-102N3) to create Pav::GFP. This BAC contains the entire *pav* locus on a 21 kb genomic fragment (chr3L:4,229,286…4,250,505, FlyBase release 6). Briefly, we used a 2-step BAC recombineering protocol to first insert a Kanamycin resistance cassette (Wang et al., 2006) which was subsequently replaced by HA::GFP::FLAG through streptomycin selection. The final plasmid was injected into BL#24872 into the attP-3B site on chr2L at Rainbow Transgenic Flies, Inc. (Camarillo, CA).

### Immunocytochemistry

Testes were dissected in IMADS buffer and fixed for 10 min in 4% paraformaldehyde in PBT (phosphate-buffered saline with 0.3% Triton X-100 and 0.5% BSA). Fixed tissue was washed in PBT and incubated with anti-Hts1B1 (1:50, Developmental Studies Hybridoma Bank (DSHB), Zaccai and Lipshitz (1996)) or anti-*α*Spec (1:50, DSHB, Dubreuil et al. (1987)). Secondary antibodies used were goat anti-mouse conjugated to Alexa-568 (1:500, Invitrogen). Samples were washed in PBT and mounted on slides in Aqua PolyMount (Polysciences, Inc.). Samples were imaged with a Leica SP8 confocal microscope and a 40× 1.3 NA oil-immersion objective lens.

### Photo-activation of PA-GFP

Live testes expressing Pav::GFP and PA-GFP with or without *α-Spectrin* shRNA driven by *nos-* or *bam-Gal4* drivers were dissected in a small drop of IMADS (ionically matched Drosophila saline, Singleton and Woodruff (1994)) on a coverslip. Testes were gently scored to release both spermatogonia and spermatocytes and break the muscle to prevent muscle contraction and keep the testes from shifting during imaging. A slide was placed over the coverslip squashing the testes and extra buffer was wicked away using a Kimwipe. The slide was sealed with VALAP (equal parts vaseline, lanolin, and paraffin) and imaged within 15 minutes. Photoactivation and subsequent live imaging of PA-GFP was accomplished on an inverted Leica SP8 confocal microscope and a 40 × 1.3 NA oil-immersion objective lens using the FRAP mode. Photoactivation was accomplished with 30 scan iterations of 405 nm light over regions of interest. Activated GFP was observed by capturing a single z-slice using 488 nm excitation every 30 seconds for ∼10 minutes.

### Male fertility assay

Fertility of the fusome-less males was assessed by pairing a single male with three CantonS or *w*^*1118*^ virgin females. These pairs were shifted to new vials every two days for 14 days and total number of adult progeny was counted to determine fertility.

### Transmission electron microscopy

For analysis by EM, ∼20 testes samples were fixed in 2.5% gluteraldehyde and 2% paraformaldehyde in 0.1M sodium cacodylate buffer pH 7.4 for 30 minutes at room temperature followed by 1 hour at 4°C. The samples were rinsed in buffer then post-fixed in 1% osmium tetroxide and en bloc stained in 2% aqueous uranyl acetate for one hour. Tissue was rinsed and dehydrated in an ethanol series followed by epon resin (Embed812 Electron Microscopy Science) infiltration, oriented, and baked overnight at 60°C. Hardened blocks were cut using a Leica UltraCut UCT. 60 nm sections were collected on formvar/carbon coated grids and contrast stained using 2% uranyl acetate and lead citrate. Samples were viewed using FEI Tencai Biotwin TEM at 80Kv. Images were taken using a Morada CCD and iTEM (Olympus) software.

### Fluorescence Loss in Photobleaching (FLIP)

FLIP of UAS-GFP and all GFP-traps was conducted using a Leica SP8 microscope. 16- or 64-cell cysts were dissected out of the testis in a manner similar to above. Microscope pinhole size was set to 7 to generate visible bleaching of GFP using the following sequence: [1 pre-bleach, 30 iterations of bleaching, 1 post-bleach] × 48 for a ∼1 hour of imaging while repositioning the sample as necessary to account for drift. GFP bleaching and image capture was performed using a 488 nm laser. All images were processed using FIJI. To quantify fluorescence loss, an ROI was drawn around the entire bleached area of a cyst and the mean pixel intensity for this region was measured for the duration of the movie using Time Series Analyzer (FIJI). Similarly, an ROI was drawn around the remainder of the cyst, outside of the bleached region, and the mean pixel intensities were measured. These values were plotted against the mean pixel values from an ROI of similar size in a neighboring control cell to control for loss of signal due to photobleaching. Raw values of the mean pixel intensities were exported into GraphPad Prism to generate representative graphs of each GFP protein.

### Quantitative Analysis of GFP Movement

Movement of GFP fluorescence, both in PA-GFP and FLIP experiments, was assessed by recording fluorescence intensities in experimental and non-activated control cells over the course of the movies. Average pixel intensity values from a 1256 pixel region of interest (ROI) in the cytoplasm of either the donor or acceptor cell(s) were measured using the Time Series Analyzer in FIJI. In cysts with more than one acceptor cell, all cells in the focal plane were averaged to generate a single trace. Fluorescence values were normalized by subtracting the average fluorescence intensity of an ROI of the same size from two adjacent, non-photoactivated cells. Measurements were exported into Excel for further analysis and GraphPad Prism for data visualization.

Measurements of GFP movement in elongated spermatids were acquired using the Time Series Analyzer in FIJI. Six 1256 pixel ROIs were assigned as described in Figure 7P and the mean pixel values of each ROI were plotted as a function of time. Raw values were exported into Excel for further analysis and GraphPad Prism for data visualization. A total of three spermatid bundles were analyzed and plotted individually because of differing levels of GFP fluorescence.

To quantitatively assess rates of movement of PA-GFP between cells in wild-type and fusome knockdown testes, a Bruker Opterra II Swept Field Microscope was used, with a 60× water immersion objective lens. Wild-type (*w*^*1118*^) and *α-Spectrin* shRNA (*bam* > *α-Spectrin*^*RNAi*^) testes were scored and mounted as described above. PA-GFP was activated by a single iteration of 405 nm light in one z-plane and movies were captured in a 15-20 slice z-stack encompassing the cyst every 10 seconds for a total of 10 minutes. Maximum intensity projections were generated in FIJI, and the total fluorescence of PA-GFP in the activated and recipient cell was measured at each time point. For comparison purposes, we only quantified movies in which PA-GFP was activated within a single spermatocyte cell and diffused into one other recipient cell. We note that although the cells imaged in these experiments were from 16-cell cysts, PA-GFP diffusion usually was restricted to 1-5 other cells indicating that the tissue-scoring preparation used may have caused cell clusters to become dissociated from the rest of the cyst. The PA-GFP relative fluorescence units (RFU) between the activated and the recipient cells was summed and normalized to 1. The mean RFU of PA-GFP in the recipient cell (as a fraction of total RFU between the two cells) for wild-type and fusome knockdown conditions was plotted over time (n=9 for wild-type; n=11 for fusome knockdown). A nonlinear regression was used to fit the data to a one phase exponential association model using the following equation:

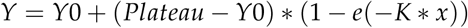

The best-fit curve was plotted alongside the mean RFU data points for each condition (see Figure 6B). Comparison of fits was performed to check for statistically significant differences in the best-fit values between wild-type and fusome knockdown fits.

## Acknowledgments

We thank Peter McLean for his help in recombineering the Pav BAC transgene; Allison James for her help screening RNAi lines; Ronit Wilk, Jack Hu, and Henry Krause for help with post-meiotic movement experiments; Julie Brill for helpful conversations about lateral perforations in spermatid tails; Tian Xu for providing access to the Leica SP8 confocal microscope; and Morven Graham and the Yale Center for Cellular and Molecular Imaging for help with EM imaging. Stocks obtained from the Bloomington Drosophila Stock Center (NIH P40OD018537) were used in this study. We thank the TRiP at Harvard Medical School for providing transgenic RNAi fly stocks used in this study.

This work was supported by the National Institutes of Health (R01 GM043301 and RC1 GM091791 to L.C.). Partial support for predoctoral trainees was provided by National Institutes of Health training grants (T32 GM007223 for R.S.K and K.M.M.).

## Author Contributions

Conceptualization: R.S.K., L.C.; Investigation: R.S.K., K.L.P., K.M.M., K.A., A.M.H., L.C., Writing – Original Draft: R.S.K.; Writing – Review & Editing: R.S.K., K.L.P., K.M.M., K.A., A.M.H., L.C.; Visualization: R.S.K., K.L.P., K.M.M.; Funding Acquisition: L.C.; Resources: R.S.K., K.L.P., K.M.M., K.A., A.M.H., L.C.; Supervision: L.C.

## Supporting Information

**Figure S1.**
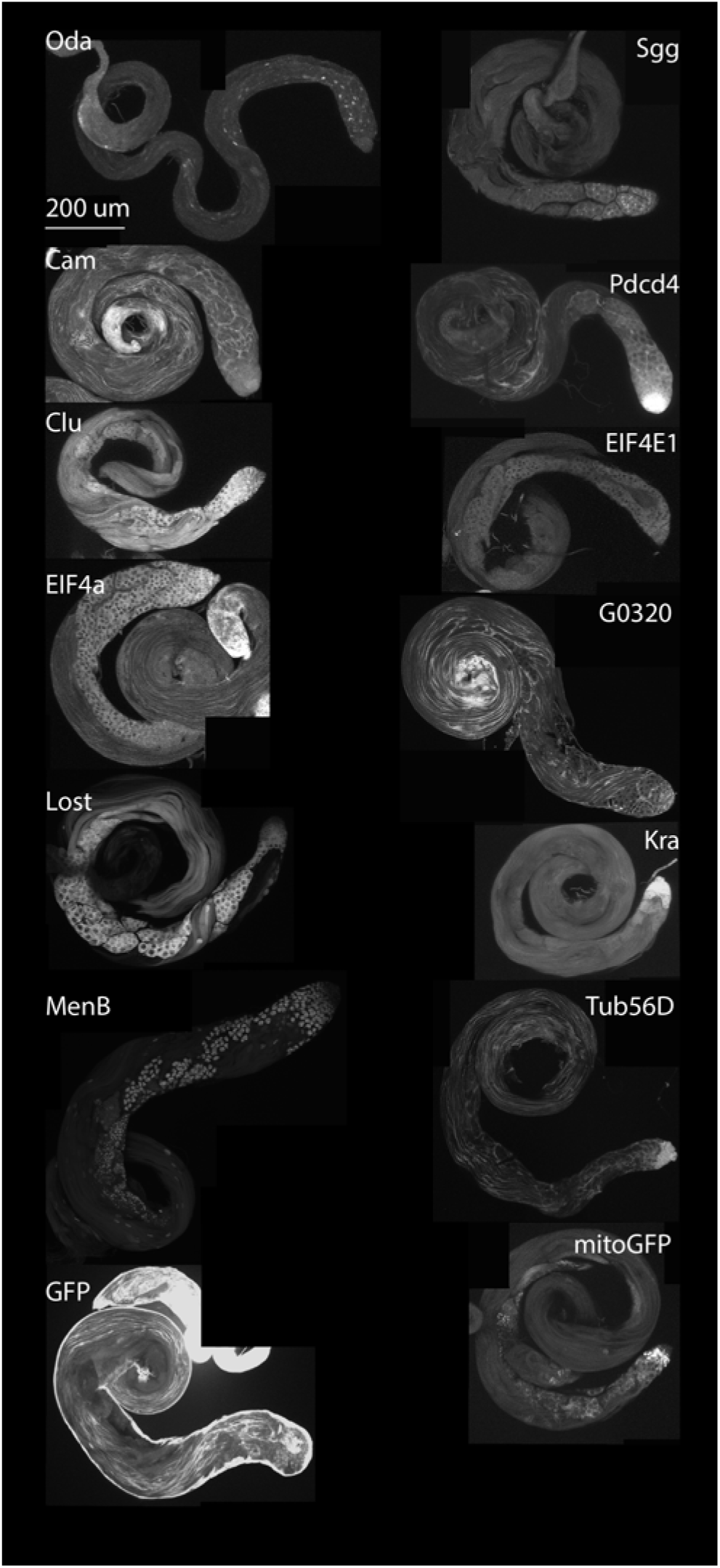
GFP expression pattern of FlyTrap lines used in FLIP imaging. Whole mount testes of each of the protein traps used for FLIP imaging to demonstrate differing patterns and levels of GFP expression.

**Figure S2.**
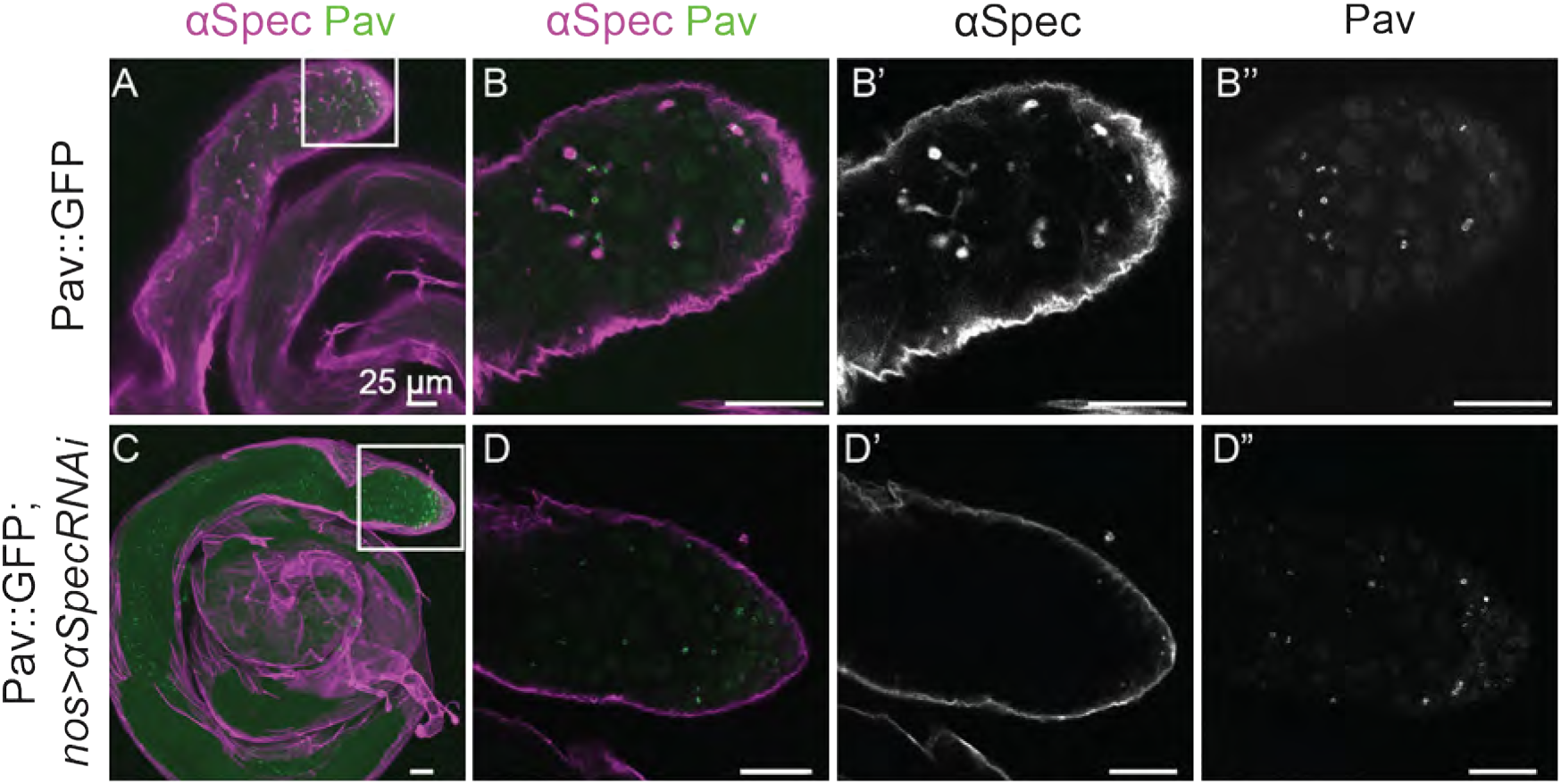
Confirming *α*-Spectrin loss using *α-Spectrin* RNAi. (A) Pav::GFP testis stained with *α*-Spectrin showed staining of the fusome similar to Hts1B1. (B-B”) Close up of region marked in (A) showing *α*-Spectrin staining at the fusome (B’) and wild-type RCs (B”). (C) Loss of fusome using *α-Spectrin* RNAi showed a lack of *α*-Spectrin staining at the fusome, but testis morphology is unaffected. (D-D”) Close up of inset from (C) showing loss of *α*-Spectrin staining (D’) but intact RCs (D”).

**Figure S3.**
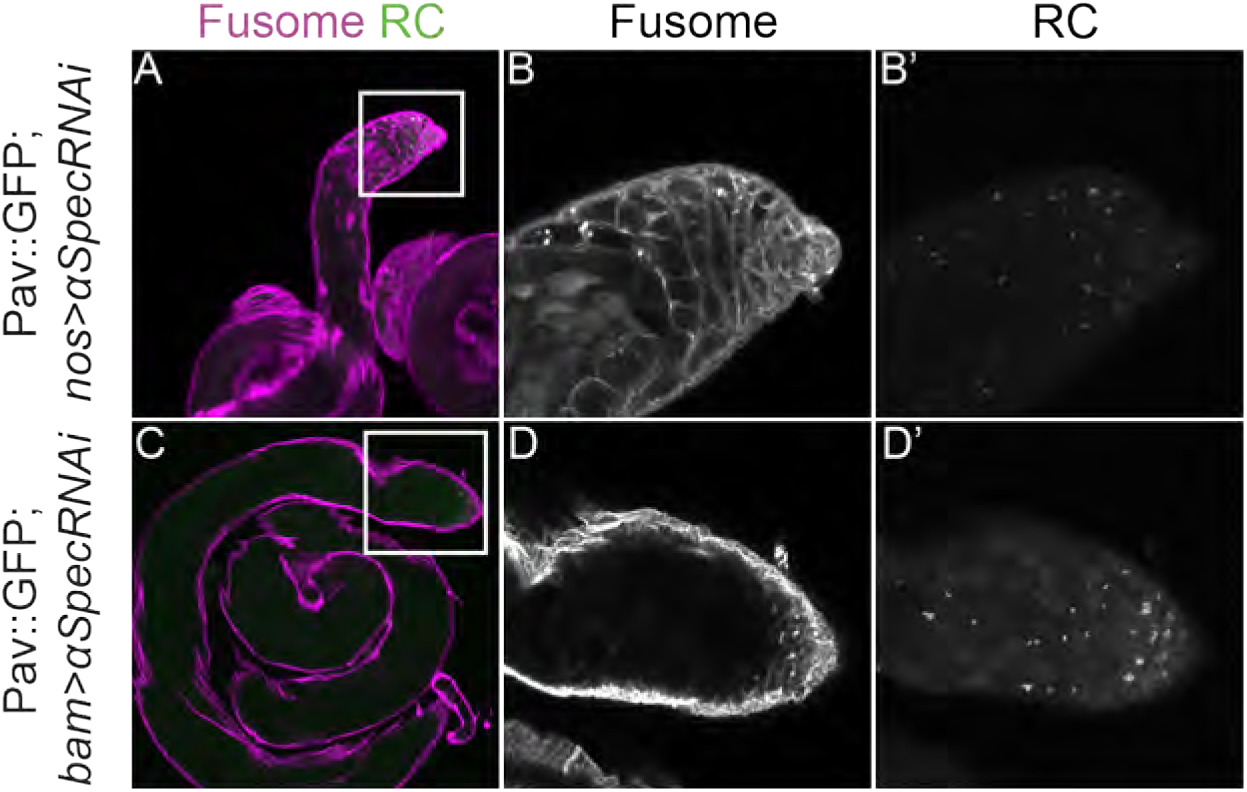
Testis morphology is unaffected by germline specific loss of the fusome. (A) Whole-mount testis of *nos-Gal4¿α-Spectrin* RNAi in tissue also expressing Pav::GFP stained with Hts1B1 antibody showed no gross abnormal morphology. (B-B’) Close up of inset in (A) showing loss of fusome (B) but intact RCs (B’). (C) Whole-mount testis of *bam-Gal4¿α-Spectrin* RNAi in tissue also expressing Pav::GFP stained with *α*-Spectrin antibody displayed no gross abnormal tissue morphology. (D-D’) Close up of inset in (C) showed loss of fusome in the post-mitotic stages (D) and normal RC localization (D’).

**Figure S4.**
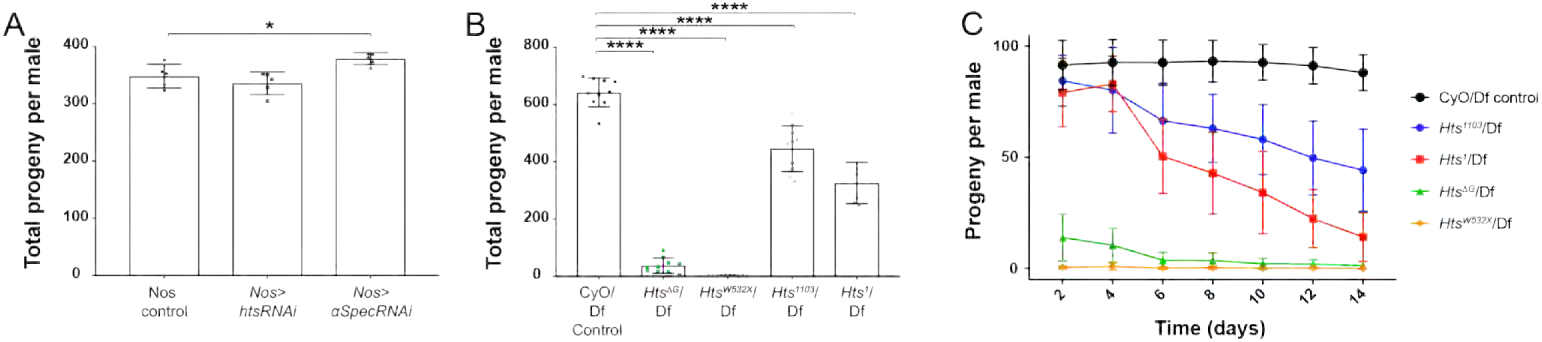
Fusome knockdown by germline specific RNAi does not have an effect on fertility. (A) Comparison of fertility as measured by total progeny per male between germline specific *nos-Gal4* control and *nos-Gal4* driving *hts* (p=0.45) or *α-Spectrin* RNAi (p=0.03) showed a slight increase in fertility in the *α-Spectrin* RNAi line. (B) Fertility assessment using *hts* alleles *hts*^Δ*G*^/Df, *hts*^*W532X*^/Df, *hts*^*1103*^/Df and *hts*^*1*^/Df showed significant decrease in fertility compared to control (p<0.0001). (C) Fertility of *hts*^*1103*^/Df and *hts*^*1*^/Df decline over time in comparison to control. *hts*^Δ*G*^/Df and *hts*^*W532X*^/Df were consistently less fertile than controls.

**Figure S5.**
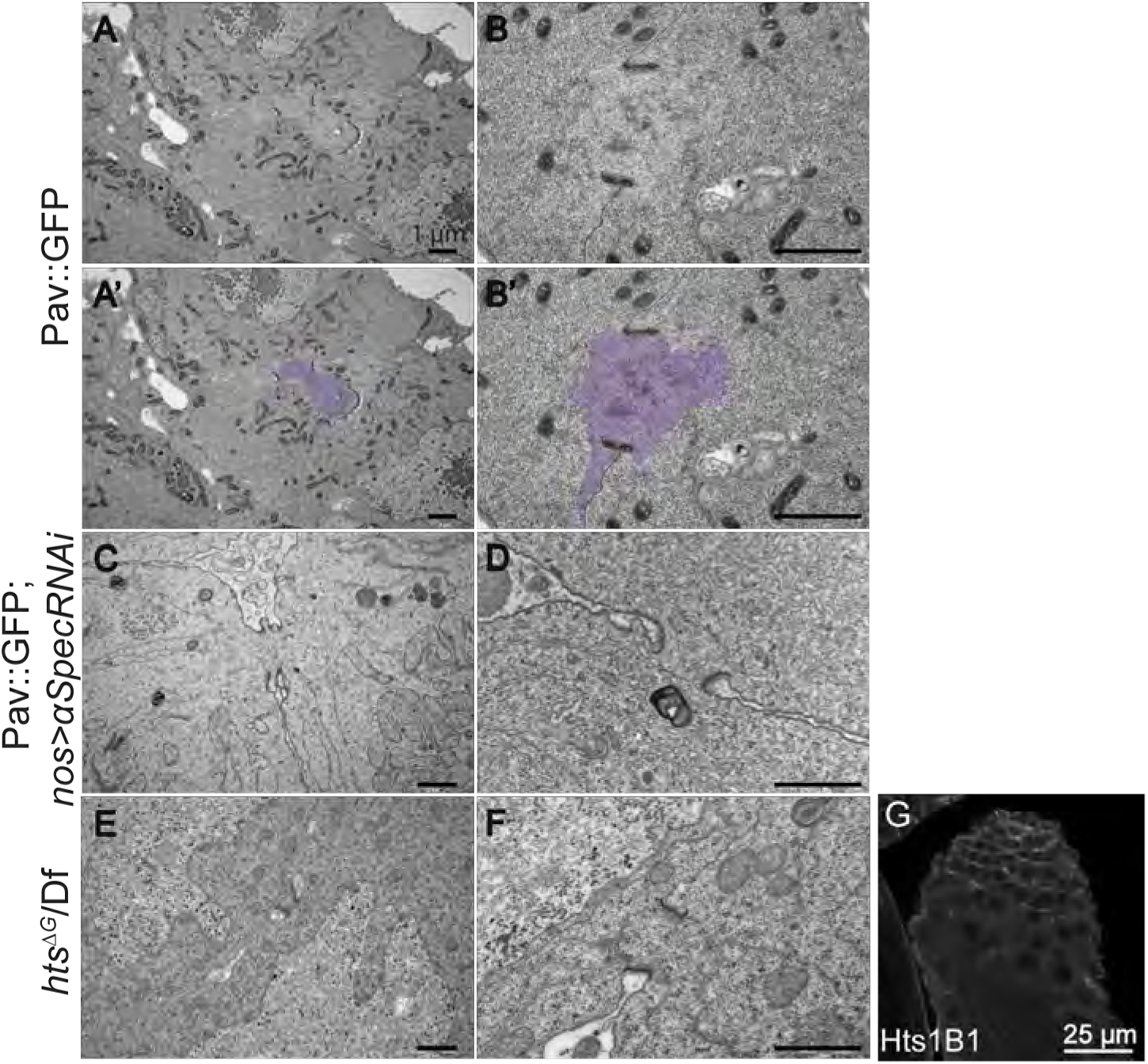
Additional EM staining showing clear fusome in wild-type and loss of fusome in *α-Spectrin* and *hts* RNAi testes. (A-B) EM of Pav::GFP testes showing RCs surrounding the fusome, a clear ribosome deficient cloud. (A’-B’) False coloring of (A-B) highlighting the fusome area in purple. (C-D) Additional EM pictures of *α-Spectrin* RNAi; showing a lack of clearly marked fusome between the electron dense RCs. (E-F) EM images of *hts*^Δ*G*^/Df testes, which similar to (C-D), show no fusome structure but intact RCs. (G) Immunofluorescence of *hts*^Δ*G*^/Df shows a lack of Hts1B1 staining at the fusome, but spermatogenesis and testis morphology appear unaffected. NOTE: A PDF FILE WAS SUBMITTED FOR VIEWING HIGH RESOLUTION OF EM IMAGES.

**Figure S6.**
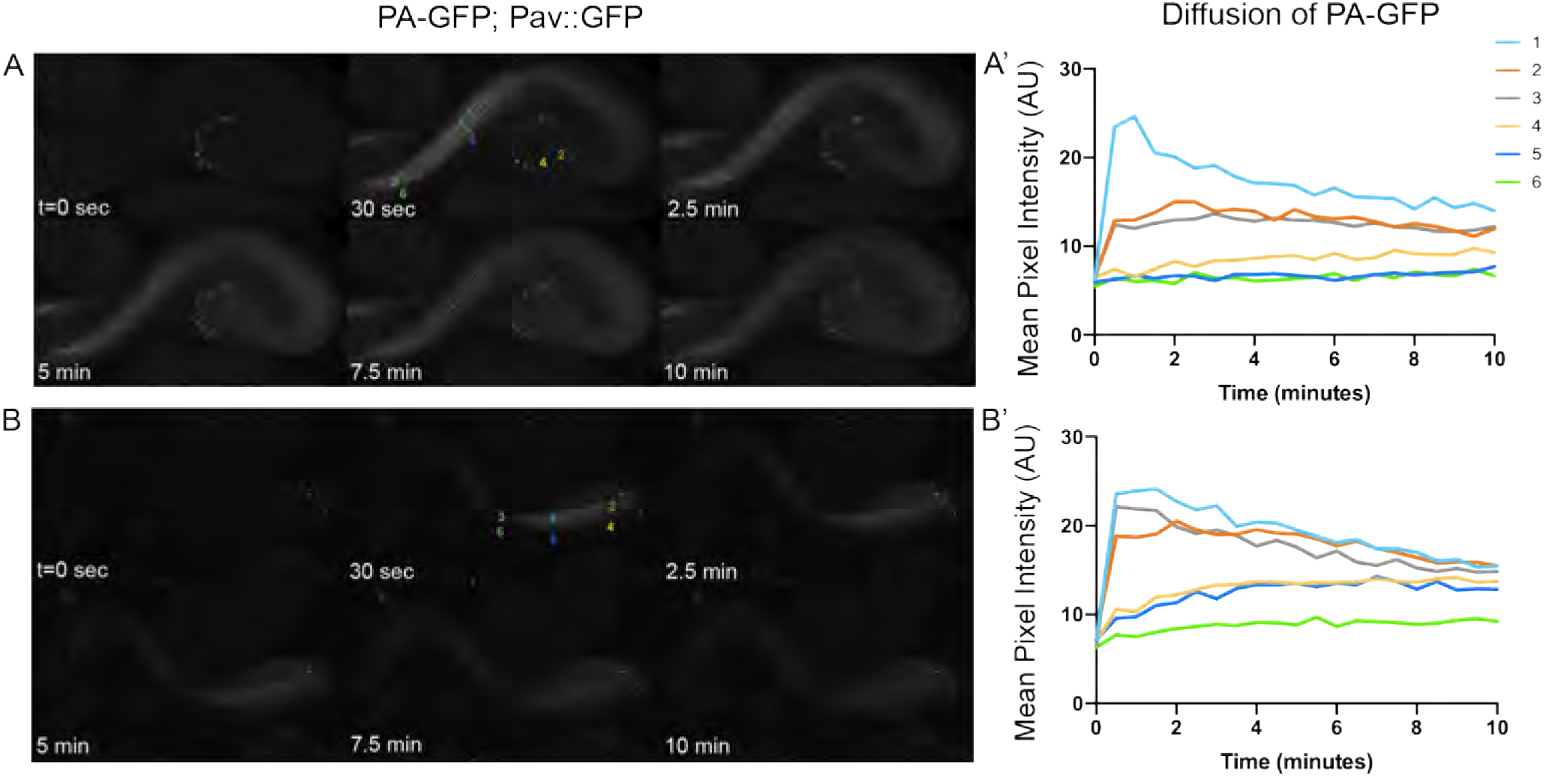
Movement of PA-GFP occurs through the RCs faster than through the tail perforations. (A-B) Still images of 10 minute movies capturing movement of PA-GFP in spermatid tails after activation in region 1. (A’-B’) Quantification of PA-GFP fluorescence in several different locations along the spermatid tails show that PA-GFP is first observed near the RCs (region 4) rather than having traveled through the perforations (region 5).

**Figure S7. Movie showing movement of PA-GFP through the RCs in a 2-cell spermatogonial cyst** Photoactivation of PA-GFP in a single cell of two 2-cell spermatogonial cysts demonstrates intercellular exchange of cytoplasmic protein. Images were acquired at 30 second intervals for 10 minutes following activation.

**Figure S8. Movie showing movement of PA-GFP through the RCs in a 4-cell spermatogonial cyst** Photoactivation of PA-GFP in a single cell of a 4-cell spermatogonial cyst demonstrates rapid intercellular exchange of GFP. Images were acquired at 30 second intervals for 10 minutes following activation.

**Figure S9. Movie showing movement of PA-GFP through the RCs in an 8-cell spermatogonial cyst** Photoactivation of PA-GFP in a single cell of an 8-cell spermatogonial cyst demonstrates intercellular exchange GFP through RCs in later stage mitotic cysts. Images were acquired at 30 second intervals for 10 minutes following activation.

**Figure S10. Movie showing movement of PA-GFP through the RCs in a 16-cell spermatocyte cyst** Photoactivation of PA-GFP in a single, center cell of a 16-cell primary spermatocyte cyst demonstrates intercellular exchange of cytoplasmic protein occurs even in cysts undergoing a growth phase. Images were acquired at 30 second intervals for 10 minutes following activation.

**Figure S11. Movie showing movement of PA-GFP through the RCs in a 32-cell cyst** Photoactivation of PA-GFP in several cells of a 32-cell cyst demonstrates intercellular exchange of cytoplasmic protein happens during meiosis. Images were acquired at 30 second intervals for 10 minutes following activation.

**Figure S12. Movie showing movement of PA-GFP through the RCs in a 64-cell cyst** Photoactivation of PA-GFP in 2 single cells of a 64-cell post-meioitc cyst demonstrates protein exchange occurs post-meiotically. Images were acquired at 30 second intervals for 10 minutes following activation.

**Figure S13. Movie showing movement of PA-GFP through the RCs in elongating spermatids** Photoactivation of PA-GFP in a subset of spermatids demonstrates intercellular exchange of cytoplasmic protein occurs post-tail elongation. Images were acquired at 30 second intervals for 10 minutes following activation.

**Figure S14. Moving showing movement of PA-GFP through the RCs in 2- and 4-cell spermatogonial cysts lacking a fusome** Photoactivation of PA-GFP in single cells of 2- and 4-cell spermatogonial cysts in *nos-Gal4* > *α-Spectrin* RNAi testes demonstrates intercellular exchange of cytoplasmic protein occurs despite the lack of fusome structure. Images were acquired at 30 second intervals for 10 minutes following activation.

**Figure S15. Movie showing movement of PA-GFP through the RCs in a 16-cell spermatocyte cyst lacking a fusome** Photoactivation of PA-GFP in a single cell of 16-cell spermatocyte cyst in *bam-Gal4* > *α-Spectrin* RNAi testes demonstrates intercellular exchange of PA-GFP is not mediated by the fusome in primary spermatocytes. Images were acquired at 30 second intervals for 10 minutes following activation.

**Figure S16. Movie showing movement of PA-GFP through the RCs in a spermatids lacking a fusome** Photoactivation of PA-GFP in a subset spermatids in *bam-Gal4* > *α-Spectrin* RNAi testes demonstrates intercellular exchange of PA-GFP is not mediated by the fusome post-meiotically. Images were acquired at 30 second intervals for 40 minutes following activation.

## References

Airoldi, S. J., P. F. McLean, Y. Shimada, and L. Cooley (2011). “Intercellular protein movement in syncytial Drosophila follicle cells”. en. Journal of Cell Science 124.23, pp. 4077–4086. DOI:10.1242/jcs.090456.

Barreau, C., E. Benson, E. Gudmannsdottir, F. Newton, and H. White-Cooper (2008). “Post-meiotic transcription in Drosophila testes”. en. Development 135.11, pp. 1897–1902. DOI:10.1242/dev.021949.

Braun, R. E., R. R. Behringer, J. J. Peschon, R. L. Brinster, and R. D. Palmiter (1989). “Genetically haploid spermatids are phenotypically diploid”. Nature 337, p. 373.

Brown, E. H. and R. C. King (1964). “Studies on the events resulting in the formation of an egg chamber in drosophila melanogaster”. eng. Growth 28, pp. 41–81.

Burgos, M. H. (1955). “Studies on the fine structure of the mammalian testis: i. differentiation of the spermatids in the cat (felis domestica)”. en. The Journal of Cell Biology 1.4, pp. 287–300. DOI:10.1083/jcb.1.4.287.

Buszczak, M., S. Paterno, D. Lighthouse, J. Bachman, J. Planck, S. Owen, A. D. Skora, T. G. Nystul, B. Ohlstein, A. Allen, J. E. Wilhelm, T. D. Murphy, R. W. Levis, E. Matunis, N. Srivali, R. A. Hoskins, and A. C. Spradling (2007). “The carnegie protein trap library: a versatile tool for Drosophila developmental studies”. eng. Genetics 175.3, pp. 1505–1531. DOI:10.1534/genetics.106.065961.

Chen, D. and D. M. McKearin (2003). “A discrete transcriptional silencer in the bam gene determines asymmetric division of the Drosophila germline stem cell”. en. Development 130.6, pp. 1159–1170. DOI:10.1242/dev.00325.

Cuevas, M. de, M. Lilly, and A. Spradling (1997). “Germline Cyst Formation in Drosophila”. Annual Review of Genetics 31.1, pp. 405–428. DOI:10.1146/annurev.genet.31.1.405.

Cuevas M. de and A. C. Spradling (1996). “Morphogenesis of the Drosophila fusome and its implications for oocyte specification”. en, p. 9.

Deng, W. and H. Lin (1997). “Spectrosomes and Fusomes Anchor Mitotic Spindles during Asymmetric Germ Cell Divisions and Facilitate the Formation of a Polarized Microtubule Array for Oocyte Specification inDrosophila”. Developmental Biology 189.1, pp. 79–94. DOI:10.1006/dbio.1997.8669.

Dubreuil, R., T. J. Byers, D. Branton, L. S. B. Goldstein, and Kiehart (1987). “Drosophilia spectrin. I. Characterization of the purified protein”. The Journal of Cell Biology 105.5, pp. 2095–2102.

Dym, M. and D. W. Fawcett (1971). “Further observations on the numbers of spermatogonia, spermatocytes, and spermatids connected by intercellular bridges in the mammalian testis”. eng. Biol. Reprod. 4.2, pp. 195–215.

Erickson, R. P. (1973). “Haploid gene expresion versus meiotic drive: the relevance of intercellular bridges during spermatogenesis”. eng. Nature New Biol. 243.128, pp. 210–212.

Fabrizio, J. J., G. Hime, S. K. Lemmon, and C. Bazinet (1998). “Genetic dissection of sperm individualization in Drosophila melanogaster”. en. Development 125.10, pp. 1833–1843.

Fawcett, D. W. (1959). “The Occurrence of Intercellular Bridges in Groups of Cells Exhibiting Synchronous Differentiation”. en. The Journal of Cell Biology 5.3, pp. 453–460. DOI:10.1083/jcb.5.3.453.

Fuller, M. T. (1993). “Spermatogenesis”. The development of Drosophila melanogaster. Cold Spring Harbor Press, pp. 71–147. ISBN: 978-0-87969-423-4.

Gärtner, S. M. K., C. Rathke, R. Renkawitz-Pohl, and S. Awe (2014). “Ex vivo culture of Drosophila pupal testis and single male germ-line cysts: dissection, imaging, and pharmacological treatment”. eng. J Vis Exp 91, p. 51868. DOI:10.3791/51868.

Greenbaum, M. P., T. Iwamori, G. M. Buchold, and M. M. Matzuk (2011). “Germ Cell Intercellular Bridges”. en. Cold Spring Harbor Perspectives in Biology 3.8, a005850–a005850. DOI:10.1101/cshperspect.a005850.

Greenbaum, M. P., L. Ma, and M. M. Matzuk (2007). “Conversion of midbodies into germ cell intercellular bridges”. en. Developmental Biology 305.2, pp. 389–396. DOI:10.1016/j.ydbio.2007.02.025.

Grieder, N. C., M. d. Cuevas, and A. C. Spradling (2000). “The fusome organizes the microtubule network during oocyte differentiation in Drosophila”. en. Development 127.19, pp. 4253–4264.

Haglund, K., I. P. Nezis, and H. Stenmark (2011). “Structure and functions of stable intercellular bridges formed by incomplete cytokinesis during development”. en. Communicative & Integrative Biology 4.1, pp. 1–9. DOI:10.4161/cib.13550.

Hime, G. R., J. A. Brill, and M. T. Fuller (1996). “Assembly of ring canals in the male germ line from structural components of the contractile ring”. en, p. 10.

Huckins, C. (1978). “Spermatogonial intercellular bridges in whole-mounted seminiferous tubules from normal and irradiated rodent testes”. en. American Journal of Anatomy 153.1, pp. 97–121. DOI:10.1002/aja.1001530107.

Huynh, J.-R. (2013). Fusome as a Cell-Cell Communication Channel of Drosophila Ovarian Cyst. en. Landes Bioscience.

Koch, E. A. and R. C. King (1969). “Further studies on the ring canal system of the ovarian cystocytes of Drosophila melanogaster”. eng. Z Zellforsch Mikrosk Anat 102.1, pp. 129–152.

Koch, E. A. and R. C. King (1966). “The origin and early differentiation of the egg chamber of Drosophila melanogaster”. en. Journal of Morphology 119.3, pp. 283–303. DOI:10.1002/jmor.1051190303.

Koundakjian, E. J., D. M. Cowan, R. W. Hardy, and A. H. Becker (2004). “The Zuker Collection: A Resource for the Analysis of Autosomal Gene Function in Drosophila melanogaster”. en. Genetics 167.1, pp. 203–206. DOI:10.1534/genetics.167.1.203.

LeGrand, E. K. (2001). “Genetic conflict and apoptosis”. eng. Perspect. Biol. Med. 44.4, pp. 509–521.

Lei, L. and A. C. Spradling (2016). “Mouse oocytes differentiate through organelle enrichment from sister cyst germ cells”. en. Science 352.6281, pp. 95–99. DOI:10.1126/science.aad2156.

Lighthouse, D. V., M. Buszczak, and A. C. Spradling (2008). “New components of the Drosophila fusome suggest it plays novel roles in signaling and transport”. en. Developmental Biology 317.1, pp. 59–71. DOI:10.1016/j.ydbio.2008.02.009.

Lilly, M. A., M. de Cuevas, and A. C. Spradling (2000). “Cyclin A Associates with the Fusome during Germline Cyst Formation in the Drosophila Ovary”. en. Developmental Biology 218.1, pp. 53–63. DOI:10.1006/dbio.1999.9570.

Lin, H., L. Yue, and A. C. Spradling (1994). “The Drosophila fusome, a germline-specific organelle, contains membrane skeletal proteins and functions in cyst formation”. en. Development 120.4, pp. 947–956.

Lin, H. and A. C. Spradling (1995). “Fusome asymmetry and oocyte determination in Drosophila”. en. Developmental Genetics 16.1, pp. 6–12. DOI:10.1002/dvg.1020160104.

Lindsley, D. L. and E. H. Grell (1969). “Spermiogenesis without chromosomes in Drosophila melanogaster”. eng. Genetics 61.1, Suppl:69–78.

Lindsley, D. T. and K. T. Tokuyasu (1978). “Spermatogenesis”. The Genetics and Biology of Drosophila. New York, pp. 225–294.

Lowe, N., J. S. Rees, J. Roote, E. Ryder, I. M. Armean, G. Johnson, E. Drummond, H. Spriggs, J. Drummond, J. P. Magbanua, H. Naylor, B. Sanson, R. Bastock, S. Huelsmann, V. Trovisco, M. Landgraf, S. Knowles-Barley, J. D. Armstrong, H. White-Cooper, C. Hansen, R. G. Phillips, The UK Drosophila Protein Trap Screening Consortium, K. S. Lilley, S. Russell, and D. St Johnston (2014). “Analysis of the expression patterns, subcellular localisations and interaction partners of Drosophila proteins using a pigP protein trap library”. en. Development 141.20, pp. 3994–4005. DOI:10.1242/dev.111054.

Lu, K. L. and Y. M. Yamashita (2017). “Germ cell connectivity enhances cell death in response to DNA damage in the Drosophila testis”. eLife 6. Ed. by H. J. Bellen, e27960. DOI:10.7554/eLife.27960.

McGrail, M. and T. S. Hays (1997). “The microtubule motor cytoplasmic dynein is required for spindle orientation during germline cell divisions and oocyte differentiation in Drosophila”. en. Development 124.12, pp. 2409–2419.

McLean, P. F. and L. Cooley (2013). “Protein Equilibration Through Somatic Ring Canals in Drosophila”. en. Science 340.6139, pp. 1445–1447. DOI:10.1126/science.1234887.

Miyauchi, C., D. Kitazawa, I. Ando, D. Hayashi, and Y. H. Inoue (2013). “Orbit/CLASP Is Required for Germline Cyst Formation through Its Developmental Control of Fusomes and Ring Canals in Drosophila Males”. en. PLoS ONE 8.3. Ed. by C. Prigent, e58220. DOI:10.1371/journal.pone.0058220.

Morales, C. R., S. Lefrancois, V. Chennathukuzhi, M. El-Alfy, X. Wu, J. Yang, G. L. Gerton, and N. B. Hecht (2002). “A TB-RBP and Ter ATPase Complex Accompanies Specific mRNAs from Nuclei through the Nuclear Pores and into Intercellular Bridges in Mouse Male Germ Cells”. Developmental Biology 246.2, pp. 480–494. DOI:10.1006/dbio.2002.0679.

Nagarkar-Jaiswal, S., P.-T. Lee, M. E. Campbell, K. Chen, S. Anguiano-Zarate, M. C. Gutierrez, T. Busby, W.-W. Lin, Y. He, K. L. Schulze, B. W. Booth, M. Evans-Holm, K. J. Venken, R. W. Levis, A. C. Spradling, R. A. Hoskins, and H. J. Bellen (2015). “A library of MiMICs allows tagging of genes and reversible, spatial and temporal knockdown of proteins in Drosophila”. en. eLife Sciences 4, e05338. DOI:10.7554/eLife.05338.

Pfeiffer, B. D., J. W. Truman, and G. M. Rubin (2012). “Using translational enhancers to increase transgene expression in Drosophila”. en. Proceedings of the National Academy of Sciences 109.17, pp. 6626–6631. DOI:10.1073/pnas.1204520109.

Quiñones-Coello, A. T., L. N. Petrella, K. Ayers, A. Melillo, S. Mazzalupo, A. M. Hudson, S. Wang, C. Castiblanco, M. Buszczak, R. A. Hoskins, and L. Cooley (2007). “Exploring strategies for protein trapping in Drosophila”. eng. Genetics 175.3, pp. 1089–1104. DOI:10.1534/genetics.106.065995.

Ren, H. P. and L. D. Russell (1991). “Clonal development of interconnected germ cells in the rat and its relationship to the segmental and subsegmental organization of spermatogenesis”. eng. Am. J. Anat. 192.2, pp. 121–128. DOI:10.1002/aja.1001920203.

Robinson, D. N. and L. Cooley (1996). “Stable intercellular bridges in development: the cytoskeleton lining the tunnel”. en. Trends in Cell Biology 6.12, pp. 474–479. DOI:10.1016/0962-8924(96)84945-2.

Sanabria, H., M. A. Digman, E. Gratton, and M. N. Waxham (2008). “Spatial Diffusivity and Availability of Intracellular Calmodulin”. Biophysical Journal 95.12, pp. 6002–6015. DOI:10.1529/biophysj.108.138974.

Singleton, K. and R. I. Woodruff (1994). “The Osmolarity of Adult Drosophila Hemolymph and Its Effect on Oocyte-Nurse Cell Electrical Polarity”. Developmental Biology 161.1, pp. 154–167. DOI:10.1006/dbio.1994.1017.

Snapp, E. L., T. Iida, D. Frescas, J. Lippincott-Schwartz, and M. A. Lilly (2004). “The Fusome Mediates Intercellular Endoplasmic Reticulum Connectivity in Drosophila Ovarian Cysts”. Molecular Biology of the Cell 15.10, pp. 4512–4521. DOI:10.1091/mbc.E04-06-0475.

Tokuyasu, K. T., W. J. Peacock, and R. W. Hardy (1972). “Dynamics of spermiogenesis in Drosophila melanogaster. I. Individualization process”. eng. Z Zellforsch Mikrosk Anat 124.4, pp. 479–506.

Ventelä, S., J. Toppari, and M. Parvinen (2003). “Intercellular Organelle Traffic through Cytoplasmic Bridges in Early Spermatids of the Rat: Mechanisms of Haploid Gene Product Sharing”. MBoC 14.7, pp. 2768–2780. DOI:10.1091/mbc.e02-10-0647.

Wang, J., M. Sarov, J. Rientjes, J. Fu, H. Hollak, H. Kranz, W. Xie, A. F. Stewart, and Y. Zhang (2006). “An Improved Recombineering Approach by Adding RecA to λ Red Recombination”. en. Molecular Biotechnology 32.1, pp. 043–054. DOI:10.1385/MB:32:1:043.

White-Cooper, H. (2010). “Molecular mechanisms of gene regulation during Drosophila spermatogenesis”. en US. Reproduction 139.1, pp. 11–21. DOI:10.1530/REP-09-0083.

Wilk, R., J. Hu, and H. M. Krause (2017). “In Situ Hybridization: Fruit Fly Embryos and Tissues”. en. Current Protocols Essential Laboratory Techniques 15.1, pp. 9.3.1–9.3.26. DOI:10.1002/cpet.14.

Wilson, P. G. (2005). “Centrosome inheritance in the male germ line of Drosophila requires hu-li tai-shao function”. en. Cell Biology International 29.5, pp. 360–369. DOI:10.1016/j.cellbi.2005.03.002.

Yamashita, Y. M. (2018). “Subcellular Specialization and Organelle Behavior in Germ Cells”. en. Genetics 208.1, pp. 19–51. DOI:10.1534/genetics.117.300184.

Yan, D., R. A. Neumüller, M. Buckner, K. Ayers, H. Li, Y. Hu, D. Yang-Zhou, L. Pan, X. Wang, C. Kelley, A. Vinayagam, R. Binari, S. Randklev, L. A. Perkins, T. Xie, L. Cooley, and N. Perrimon (2014). “A regulatory network of Drosophila germline stem cell self-renewal”. Developmental cell 28.4, pp. 459–473. DOI:10.1016/j.devcel.2014.01.020.

Yue, L. and A. C. Spradling (1992). “hu-H tai shao, a gene required for ring canal formation during Drosophila oogenesis, encodes a homolog of adducin”. en, p. 13.

Zaccai, M. and H. D. Lipshitz (1996). “Differential distributions of two adducin-like protein isoforms in the Drosophila ovary and early embryo”. eng. Zygote 4.2, pp. 159–166.

